# Compensation between FOXP transcription factors maintains proper striatal function

**DOI:** 10.1101/2023.06.26.546567

**Authors:** Newaz I. Ahmed, Nitin Khandelwal, Ashley G. Anderson, Ashwinikumar Kulkarni, Jay Gibson, Genevieve Konopka

**Affiliations:** Department of Neuroscience, UT Southwestern Medical Center, Dallas, TX 75390-9111, USA; Peter O’Donnell Jr. Brain Institute, UT Southwestern Medical Center, Dallas, TX 75390-9111, USA; Department of Molecular and Human Genetics, Baylor College of Medicine, Houston, TX, 77030, USA; Jan and Dan Duncan Neurological Research Institute at Texas Children’s Hospital, Houston, TX, 77030, USA

## Abstract

Spiny projection neurons (SPNs) of the striatum are critical in integrating neurochemical information to coordinate motor and reward-based behavior. Mutations in the regulatory transcription factors expressed in SPNs can result in neurodevelopmental disorders (NDDs). Paralogous transcription factors *Foxp1* and *Foxp2*, which are both expressed in the dopamine receptor 1 (D1) expressing SPNs, are known to have variants implicated in NDDs. Utilizing mice with a D1-SPN specific loss of *Foxp1*, *Foxp2*, or both and a combination of behavior, electrophysiology, and cell-type specific genomic analysis, loss of both genes results in impaired motor and social behavior as well as increased firing of the D1-SPNs. Differential gene expression analysis implicates genes involved in autism risk, electrophysiological properties, and neuronal development and function. Viral mediated re-expression of *Foxp1* into the double knockouts was sufficient to restore electrophysiological and behavioral deficits. These data indicate complementary roles between *Foxp1* and *Foxp2* in the D1-SPNs.

## Introduction

The GABAergic spiny projection neurons (SPNs) of the striatum serve as a hub for important neurochemical messages from different regions of the brain. These inhibitory neurons receive dopaminergic input from the midbrain, particularly the substantia nigra (SN) and ventral tegmental area (VTA), as well as glutamatergic inputs from the thalamus and cortex. The SPNs are the primary cell-type in the striatum, the input nucleus of the basal ganglia, and play a key role in the ability of the striatum to coordinate motor activity and reward-based behavior. There are two main classes of SPNs: the dopamine receptor 1 (D1) expressing SPNs that control the direct pathway and the dopamine receptor 2 (D2) expressing SPNs that mediate the indirect pathway.^1–6^ Dysregulation of striatal circuits is implicated in many neurodevelopmental disorders (NDDs) such as autism spectrum disorder (ASD), attention deficit hyperactive disorder (ADHD), and Huntington’s disease (HD), among others.^4, 5, 7–12^ To understand the molecular mechanisms underlying these and other diseases of the striatum, it is crucial to study the factors that regulate the proper development and function of the SPNs.

Transcription factors play an essential role in the regulation of gene expression patterns which underlie the specific development, structure, and behavior of individual cell-types. This includes the striatum, where transcription factors play a key role in the differentiation, migration, and survival of striatal neurons.^13^ Since these regulatory proteins are key to normal developmental functions, genetic variants in transcription factors result in high risk for disease susceptibility.^14, 15^ The forkhead box transcription factors (Fox) are a large family of transcription factors that share a common DNA binding domain. Two members of this family, FOXP1 and FOXP2, have enriched striatal gene expression and mutations in these genes have been implicated in striatal function.^16–22^ The members of the Fox family must dimerize to function properly; this may either be in the form of homodimers or heterodimers between FOXP1 and FOXP2.^23–27^ It is thought that transcription factor dimerization arose to reduce the evolutionary constraint on the DNA binding motif of a given protein, thereby allowing for greater flexibility in transcriptional regulation.^28, 29^ Interestingly, previous studies have found that the FOXP proteins can cooperatively function to maintain proper cell function.^30–33^ However, a cooperative role for these proteins in the brain has yet to be described.

*Foxp1* and *Foxp2* are both expressed in the D1 SPNs, one of the few neuronal cell-types with high co-expression of these genes.^16–18, 20, 22, 34^ In contrast, D2 SPNs primarily express *Foxp1*.^18, 20^ Foxp1 is crucial for the development and function of the SPNs.^18, 19, 21, 35^ Foxp2 has also been linked to striatal function.^36–40^ Furthermore, genetic variants in either *FOXP1* or *FOXP2* are among the most significant recurrent *de novo* mutations associated with NDDs. Variants in *FOXP1* rank among the top genes implicated in ASD.^34, 41–44^ Variants in both genes are also associated with ASD relevant phenotypes and speech and language deficits.^21, 22, 34, 41, 45–60^ Both genes have also been implicated in attention deficit hyperactivity disorder.^43, 61–65^

We previously reported that loss of *Foxp1* from the D2-SPNs results in reduced D2 specification, impaired motor learning, hypoactivity, D2-SPN hyperexcitability, and loss of D2-SPN striatal projections. ^18, 19^ Conversely, many of these deficits are not found in the D1-SPNs upon loss of *Foxp1* except for hyperexcitability and impairments in social behavior.^18, 19, 21^ Given the specific enrichment of *Foxp2* in the D1-SPNs, this raises the possibility that Foxp2 compensates for the loss of Foxp1. In addition, striatal-specific loss of *Foxp2* also showed no major motor learning impairments, indicating reciprocal compensation by *Foxp1*.^66^ Previous work in non-neuronal tissue has investigated the synergy between *Foxp1*, *Foxp2*, and the closely related *Foxp4* where studies have found differing levels of cooperativity and redundancy between the transcription factors, depending on the cell-type.^30–33^ In addition, the cooperative role of *Foxp2* and *Foxp4* in neuroepithelial maturation has been documented.^67^ However, to our knowledge, there have been no reports on the simultaneous loss of *Foxp1* and *Foxp2* from the same cell-type in the brain. Given the co-expression of the two genes in D1-SPNs, we sought to uncover a role for functional cooperativity.

Using a cell-autonomous strategy to delete *Foxp1*, *Foxp2*, or both from D1-SPNs, we examined behavioral, electrophysiological, and transcriptomic differences in mice. In comparison to double knockout mice, the single knockouts had less severe alterations in all three categories. Thus, we find evidence of compensation between the two transcription factors to maintain striatal function due to complementary roles between *Foxp1* and *Foxp2*.

## Results

### Generation of mice with Drd1 specific loss of *Foxp1* and/or *Foxp2*

To investigate Foxp1 and Foxp2 function in D1-SPNs, we generated conditional knockout (cKO) mice of *Foxp1, Foxp2,* or both in neurons using Cre driven by the *Drd1* promoter (see Methods). Forkhead protein nomenclature follows established guidelines.^68^ We generated the following mice: *Drd1cre/+*; *Foxp1^flox/flox^* (Foxp1^D1^, deletion of Foxp1 from the D1- SPNs), *Drd1cre/+*; *Foxp2^flox/flox^* (Foxp2^D1^, deletion of Foxp2 from the D1-SPNs), *Drd1cre/+*;*Foxp1^flox/flox^; Foxp2^flox/flox^* (Foxp1/2^D1^, deletion of both Foxp1 and Foxp2 from the D1-SPNs) and Control (mice with floxed genes but no Cre); (Figure 1). Since *Drd1.Cre* is expressed starting at embryonic day 14.5 (E14.5)^18^, *Foxp1* and *Foxp2* are deleted early in striatal development.

**Figure 1:**
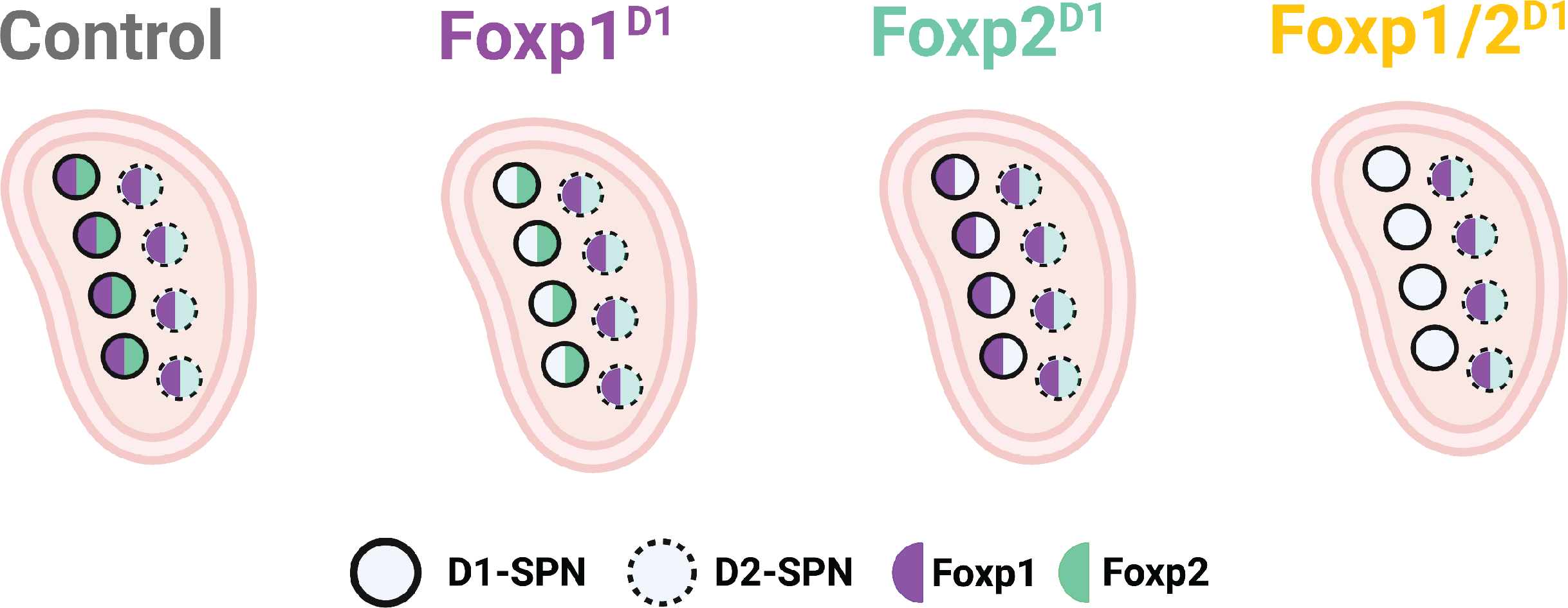
Schematic showing the genotypes used in this study. D1-cre mediated loss of *Foxp1* (Foxp1^D1^; purple), *Foxp2* (Foxp2^D1^; cyan), or both (Foxp1/2^D1^; gold), as well as cre-negative controls (grey).

### snRNA-seq of Drd1 SPNs in juvenile mice with loss of *Foxp1* and/or *Foxp2*

We had previously determined the cell-type specific transcriptional targets of Foxp1 in the juvenile striatum but hypothesized that there may be compensation by Foxp2.^18^ Therefore, we carried out snRNA-seq experiments in striatal tissue with loss of *Foxp1*, *Foxp2*, or both in the mouse brain at postnatal day 9 (P9). We profiled tissue from 3 mice from each genotype. After quality control filtering, we compared 190,082 nuclei across all four genotypes (Figures 2A and 2B). We found 11 major cell-types represented across clusters which were distributed largely equally across genotypes (Figures 2C-2E). SPNs were then subset and re-clustered to determine the contribution of each genotype to major SPN cell-types (D1, D2, and eSPNs; Figures 3A and 2F).^69^ These subsetted clusters were also filtered for quality using the same cutoffs as before retaining 52,124 D1-SPNs for further analysis (Figures 2A and 2G).

**Figure 2:**
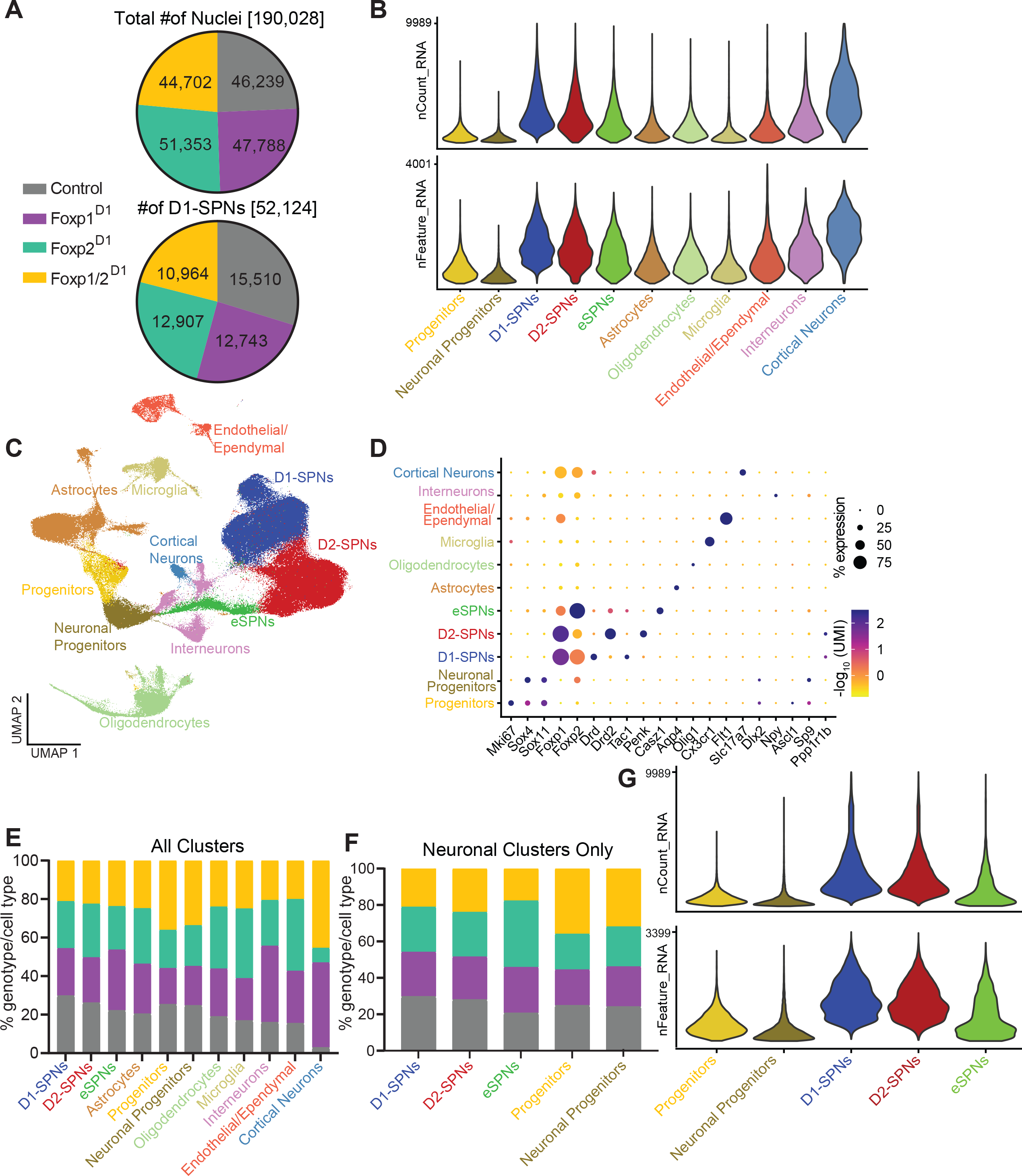
Quality control of nuclei sequenced from adult snRNA-Seq. **(A**) Pie charts showing the total number of nuclei sequenced from each genotype as well as the number of D1-SPNs profiled for each condition. **(B)** Violin plots showing the number of genes and UMI for each cell-type annotated within the dataset. **(C)** UMAP showing all of the annotated clusters from the dataset. A total of 11 major cell-types were identified. SPNs and progenitors were subset for further analysis. **(D)** Bubble plot showing overlap with marker genes used to annotate clusters. **(E)** Stacked bar plot showing the proportion of each genotype in all the cell-types as well as within the neuron-only subset **(F)** that was later generated. **(G)** Violin plots showing the number of genes and UMI for the remaining neuronal sub-types in the subset dataset.

**Figure 3:**
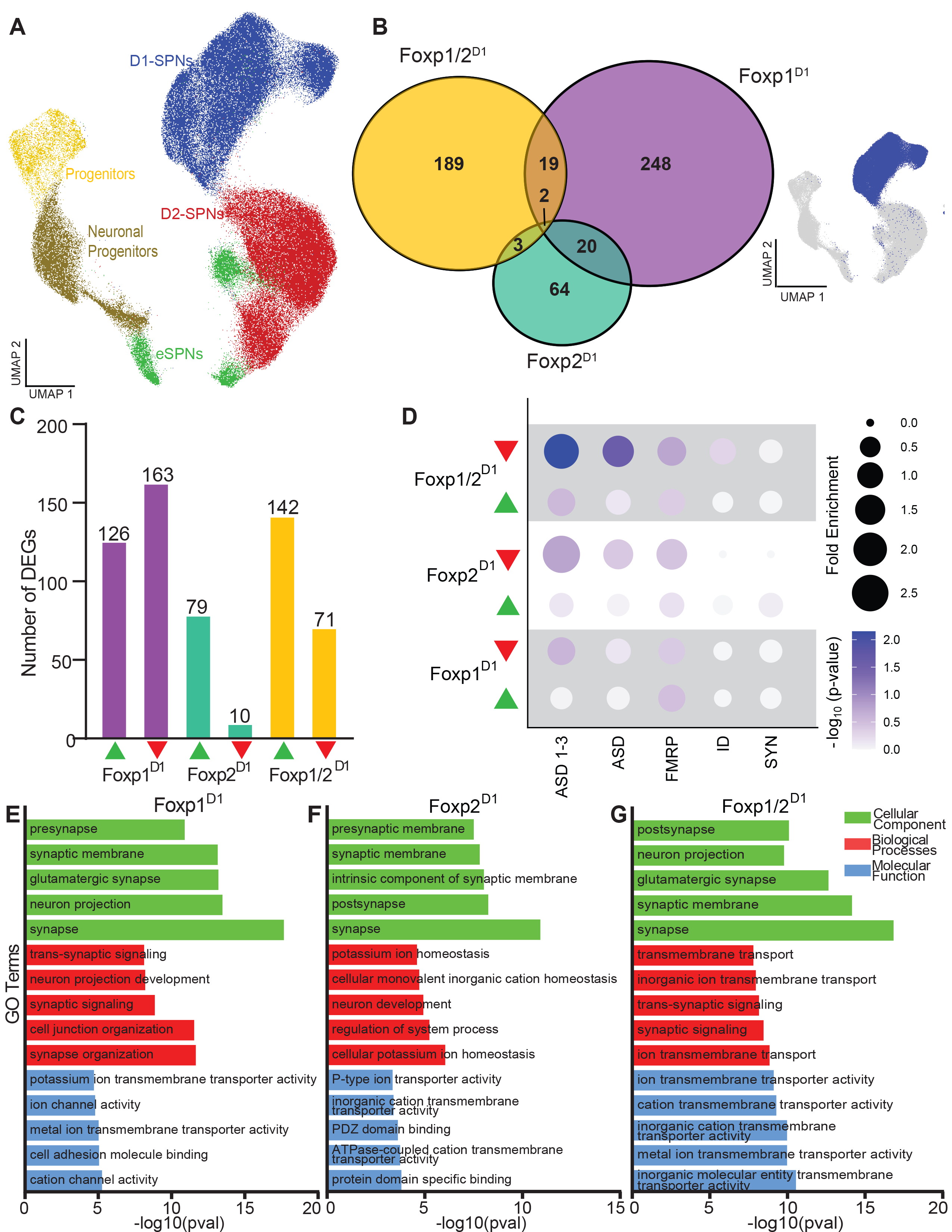
Loss of both *Foxp1* and *Foxp2* results in amplified loss of transcriptional regulation in D1-SPNs in juvenile mice. **(A**) UMAP plot generated from the neuron-only subset with colors indicating the different annotated cell-types. D1-SPNs were used for further DEG analysis. Genes were determined to be differentially expressed from controls if they had an adjusted p-value < 0.05 and absolute logFC >|0.25|. **(B)** Semi-scaled Venn diagram showing number of unique and overlapping DEGs in each knockout condition in the D1-SPNs. **(C)** Bar plots showing the number of up- and downregulated genes in each knockout condition. **(D)** Bubble chart showing enrichment of DEGs from each knockout condition. The - log10(p-value) for each enrichment is also indicated. ASD, SFARI ASD risk genes; ASD 1-3, SFARI ASD risk genes with scores of 1-3; FMRP, Fragile X Syndrome; ID, Intellectual disability; SYN, synaptic genes. Gene Ontology (GO) analysis of **(E)** Foxp1^D1^, **(F)** Foxp2^D1^, and **(G)** Foxp1/2^D1^ DEGs reveals enrichment for terms associated with electrophysiological properties and synaptic properties.

### Differentially expressed genes in juvenile D1-SPNs reveal compensation between Foxp1 and Foxp2

We next determined the differentially expressed genes (DEGs) within each cell-type. Each knockout condition was compared to controls and genes were determined to be differentially expressed based on cutoffs that we have previously used (FDR < 0.05 and an absolute log fold change of greater than 0.25).^18^ We observed 289 genes changing in the Foxp1^D1^ (126 up, 163 down), 89 in the Foxp2^D1^ (79 up, 10 down), and 213 in the Foxp1/2^D1^ (142 up, 71 down) D1-SPNs (Figures 3A and 3B). There was limited overlap between any of the conditions. Of the 213 Foxp1/2^D1^ D1-SPN DEGs, 189 (89%) were unique to that condition. These genes should represent those with *Foxp1* and *Foxp2* compensatory regulation as differential expression is only observed upon the loss of both genes. Among these 213 genes, two-thirds are upregulated, in line with previously noted repressive roles of these transcription factors (Figure 3C)^23, 26, 34, 70^.

While the overlap of differentially expressed genes between the different knockout conditions is limited, certain comparisons indicate shared regulatory mechanisms. Of the 22 genes overlapping between the Foxp1^D1^ and Foxp2^D1^ D1-SPNs, 19 changed in the same direction (Figure 3B and Table S1). These represent genes regulated similarly by both transcription factors when one is lost. There were only two genes that overlapped among all three conditions one of which was the cAMP/cGMP associated gene Phosphodiesterase 1B *Pde1b*. This gene was upregulated in the single knockouts but downregulated in the double cKO (Table S1) which again points towards a pattern of compensation. We overlapped the juvenile DEGs with a list of high confidence genes related to ASD and found enrichment with DEGs from all three cKOs (Figure 3D and Table S1). We also carried out gene ontology (GO) analysis of the DEGs and identified enrichment of similar functional terms for each knockout, in particular, categories associated with electrophysiological properties including synaptic function (Figures 3E-3G and Table S1). Our snRNA-seq in juvenile mouse D1-SPNs reveals hundreds of genes that are regulated by both *Foxp1* and *Foxp2*. These are genes that are only differentially regulated upon the loss of both transcription factors, consistent with a pattern of compensation between the two transcription factors.

### Identification of differentially accessible chromatin regions in D1 SPNs with loss of *Foxp1* and/or *Foxp2*

To determine regulatory mechanisms underlying differential gene expression with loss of *Foxp1* and/or *Foxp2,* we carried out snATAC-seq experiments in P9 mice. After quality control filtering, we profiled 75,935 nuclei from 12 mice (3mice/genotype; Figure 4A). We annotated 36 clusters using label transfer from the juvenile snRNA data (Figure 5A). All genotypes were represented across the different cell-types (Figure 4B). We determined that the number of peaks, percentage of reads in peaks, and number of reads in peaks were largely similar across genotypes (Figures 4C-4E). We next determined the differential accessibility regions (DARs) for each knockout condition in comparison to controls in the D1-SPN clusters. Interestingly, there were far more DARs in the Foxp1/2^D1^ D1-SPNs than in any other condition. We identified 237 genes with DARs in the Foxp1^D1^, 410 in the Foxp2^D1^, and 1,732 in the Foxp1/2^D1^ condition (Figure 5B and Table S2). There was more overlap between the DARs of each condition than we found for DEGs from this same time point. We found 39 DARs shared between the single knockout conditions, 51 between the Foxp1^D1^ and Foxp1/2^D1^, 149 between Foxp2^D1^ and Foxp1/2^D1^, and 46 DARs shared among all three conditions (Figure 3B). Similar percentages of DARs were unique for single cKOs (101/237, 43%, Foxp1^D1^; 177/410, 43%, Foxp2^D1^); however, most DARs in the double cKO are unique (1486/1732; 86%). This indicates that there is compensation occurring in each single cKO as the loss of both transcription factors resulted in the observation of much greater alteration in chromatin accessibility. We next asked whether genes associated with DARs overlapped with DEGs in the double cKO mice. Among the 213 DEGs in Foxp1/2^D1^ D1-SPNs, 45 (21%) also had a DAR associated with the same gene (Figure 5C).

**Figure 4:**
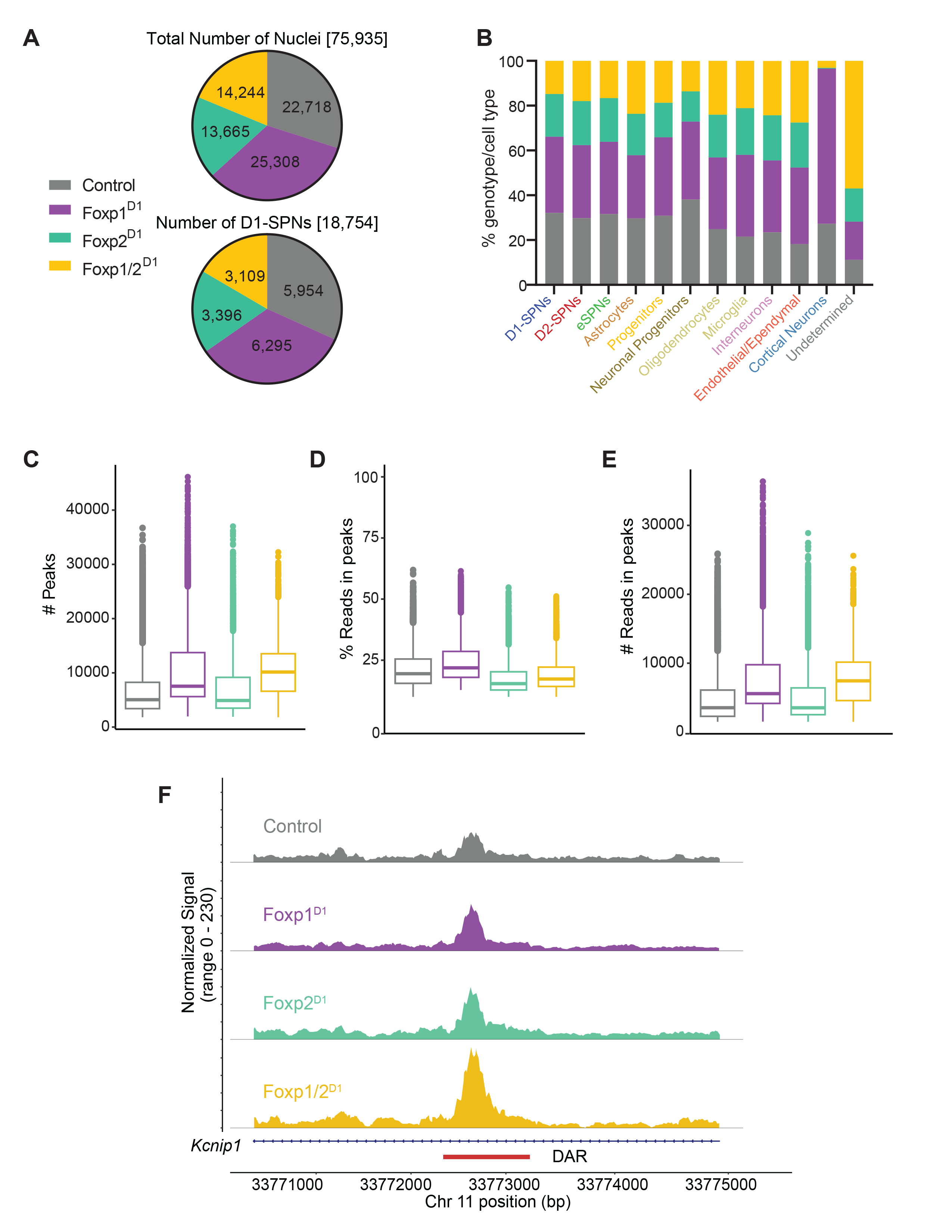
Quality control of juvenile snATAC-Seq. **(A)** Pie charts showing the total number of nuclei sequenced from each genotype as well as the number of D1-SPNs profiled for each condition. **(B)** Stacked bar plots showing the proportion of each genotype within the cell-types in the dataset. Plots showing the number of peaks (**D-C)**, percent of reads in peaks **(C)**, and number of reads in peaks **(E)** in each knockout condition. **(F)** Trackfile for *Kcnip1* with the differentially accessible region highlighted.

**Figure 5:**
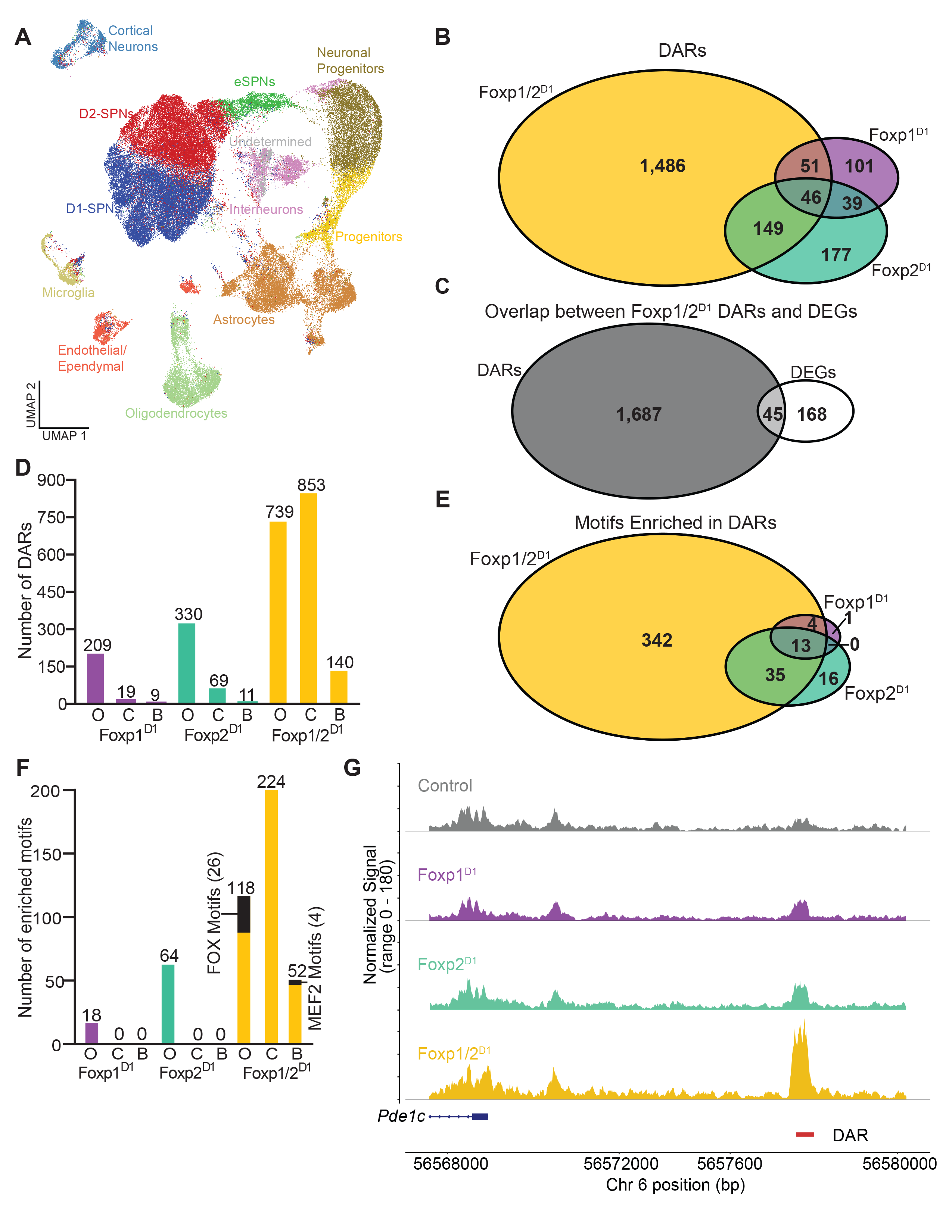
Loss of both *Foxp1* and *Foxp2* dysregulates chromatin state in D1-SPNs. **(A**) UMAP showing annotated cell-types in nuclei collected for snATAC-Seq analysis. D1-SPNs were used for further DAR analysis. A region was deemed to be differentially accessible if it had an adjusted p-value < 0.05 and a logFC > |0.1375|. **(B)** Semi-scaled Venn diagram showing the number of unique and overlapping DARs within each condition. **(C)** Venn diagrams showing the overlap between differentially expressed genes and DARs in Foxp1/2^D1^ D1-SPNs. **(D)** Bar plot showing the number of more open, more closed, or both more open and more closed regions in each knockout condition. Motifs enriched in the DARs of each knockout were identified. **(E)** Semi-scaled Venn diagram showing the number of unique and overlapping motifs in each genotype. **(F)** Bar plot showing the number of motifs enriched within more open, more closed, or both more open and more closed chromatin regions in each knockout condition. FOX and MEF2 motifs highlighted to indicate where they were enriched. **(G)** Trackfile for *Pde1c* with the differentially accessible region highlighted.

*Foxp1* and *Foxp2* genes are classically thought to be repressors.^23, 26, 34, 70^ However, changes in chromatin accessibility were roughly equivalent for more open (739) and more closed (853) DARs in the double cKOs (with some genes having both open and closed DARs associated with them). In contrast, single cKO lines had mostly open DARs with 209 open and 19 closed DARs in Foxp1^D1^ and 303 open and 69 closed DARs in Foxp2^D1^ (Figure 5D). The results in the single cKO mice support a repressive role for these transcription factors but upon the loss of both factors there are likely indirect mechanisms also at play. Consistent with our snRNA-seq findings, we again report that loss of both *Foxp1* and *Foxp2* results in large-scale dysregulation of the chromatin regions in the D1-SPNs.

### FOXP motif enrichment among DARs

We next investigated if the regions regulated by the Foxp transcription factors were direct targets or the result of further downstream regulation. Methods to determine transcription factor binding to DNA at genome-wide scale are challenging to carry out in a cell-type specific manner. Therefore, we harnessed the snATAC-seq data to determine potential direct versus indirect targets of Foxp1 and Foxp2 in D1-SPNs by assessing transcription factor motif enrichment in DARs from each cKO. Using cutoffs of FDR <0.05 and a fold enrichment of at least 1.75 within each DAR, we found enriched motifs within D1-SPNs. Foxp1^D1^ DARs had the fewest enriched motifs with only 18. Foxp2^D1^ DARs had 64 enriched motifs whereas Foxp1/2^D1^ DARs were enriched for 446 transcription factor binding motifs. Again, we found that results from Foxp1/2^D1^ cells were the most different, with these DARs containing the greatest number of unique motifs (342/446; 77%), whereas the single cKO had few unique motifs (only 1/18 for Foxp1^D1^ and 16/64 for the Foxp2^D1^; Figure 5E). Interestingly, in the single knockouts, all enriched motifs were in DARs that were more accessible whereas in the double cKOs, the motifs were enriched in both more accessible regions (118 motifs) as well as more closed DARs (224), as well as some that were associated with regions both more open and more closed (52; Figure 5F). This is consistent with the patterns observed in the DARs where the single cKOs primarily had differentially open chromatin whereas the double cKO had greater dysregulation resulting in both more open and more closed chromatin.

Given the presumed repressive role of Foxp1 and Foxp2, we hypothesized that there would be Fox motifs in the DARs associated with more open regions. This was indeed the case in the Foxp1/2^D1^ as 26/118 (22%) of the enriched motifs in more accessible chromatin regions were Fox motifs, including motifs for Foxp1 and Foxp2 (Figure 5F and Table S2). There were no such motifs enriched in more closed chromatin regions. We also identified 693 genes that contained a Fox motif enriched within an associated DAR in the double cKO (Table S2). This suggests that 693/739 (94%) genes with more open DARs are directly regulated by both Foxp1 and Foxp2. Examples of such genes include *Kcnip1*, the potassium voltage-gated channel interacting protein 1 and Phosphodiesterase 1C (*Pde1c*), which like *Pde1b* is involved in cAMP regulation (Figures 4F and 5G). The lack of Fox motif enrichment in genes associated with more closed DARs implies that, while loss of both genes can result in closed chromatin, the dysregulation is likely mediated through indirect mechanisms. To further support the idea that compensation is occurring, there were no enriched Fox motifs in either single knockout condition in both open and closed DARs.

Among non-Fox motifs that were enriched in DARs and potentially of interest for understanding striatal function, we noted motifs for the Myocyte Enhancer Binding Factor-2 (Mef2) family of transcription factors in both open and closed DARs in the Foxp1/2^D1^ data (Figure 5F). Members of this family of transcription factors are known to be directly repressed by Foxp1 or Foxp2. ^36, 71^ Some of the genes indirectly dysregulated could thus be attributed to dysregulation of *Mef2* genes due to loss of Foxp1 and Foxp2. In sum, we report that loss of both *Foxp1* and *Foxp2* results in a greater amount of altered chromatin state in comparison to either single knockout condition, indicating robustness between the two genes. These differentially accessible regions are associated with motifs enriched for binding by forkhead box proteins and their known targets.

### *Foxp1* and *Foxp2* synergistically mediate D1-SPN hyperexcitability

We previously reported that either a full body heterozygous deletion or a D2-SPN specific loss of *Foxp1* results in increased intrinsic excitability of the D2-SPNs.^19, 21^ These changes are at least partially driven by the downregulation of channels regulating potassium inward rectifying (KIR) currents such as the potassium voltage-gated channel subfamily J members 2 (*Kcnj2*) and 4 (*Kcnj4*) and potassium leak (KLeak) currents such as the potassium two pore domain channel subfamily K member 2 (*Kcnk2*).^18, 19^ We also observed increased excitability in the D1-SPNs in the Foxp1^D1^ mice, albeit to a lesser degree.^19^ This attenuated phenotype suggested the possibility that loss of *Foxp1* might be partially compensated for by *Foxp2*. To investigate this, we performed current clamp experiments in juvenile mice (P14-P18). Expression of Drd1-tdTomato was used to identify D1-SPNs in slices. In agreement with our previous work, Foxp1^D1^ D1-SPNs had increased excitability compared to controls as observed by plotting the number of action potentials as a function of injected current amplitude. (Figure 6A). D1-SPNS in Foxp1/2^D1^ mice were more hyperexcitable, even in comparison to the Foxp1^D1^ mice, including at lesser current injections (Figures 6A and 7A). Changes in subthreshold membrane properties likely contribute to the observed hyperexcitability as both Foxp1^D1^ and Foxp1/2^D1^ D1-SPNs show increased input resistance (Figure 6B). Conversely, Foxp2^D1^ D1-SPNs show significant hypoexcitability compared to controls (Figure 6A). Based on our previous work, we hypothesized that this hypoexcitability could be due to increased expression of *Foxp1* ^18^. Indeed, when we FACS sorted for td-Tomato positive D1-SPNs from Foxp2^D1^ mice at P14, we found increased *Foxp1* expression compared to controls via RT-qPCR (Figure 7B). Thus, while we now find that *Foxp2* also plays a role in the regulation of intrinsic excitability of SPNs, the role of *Foxp1* appears to eclipse that of *Foxp2*.

**Figure 6:**
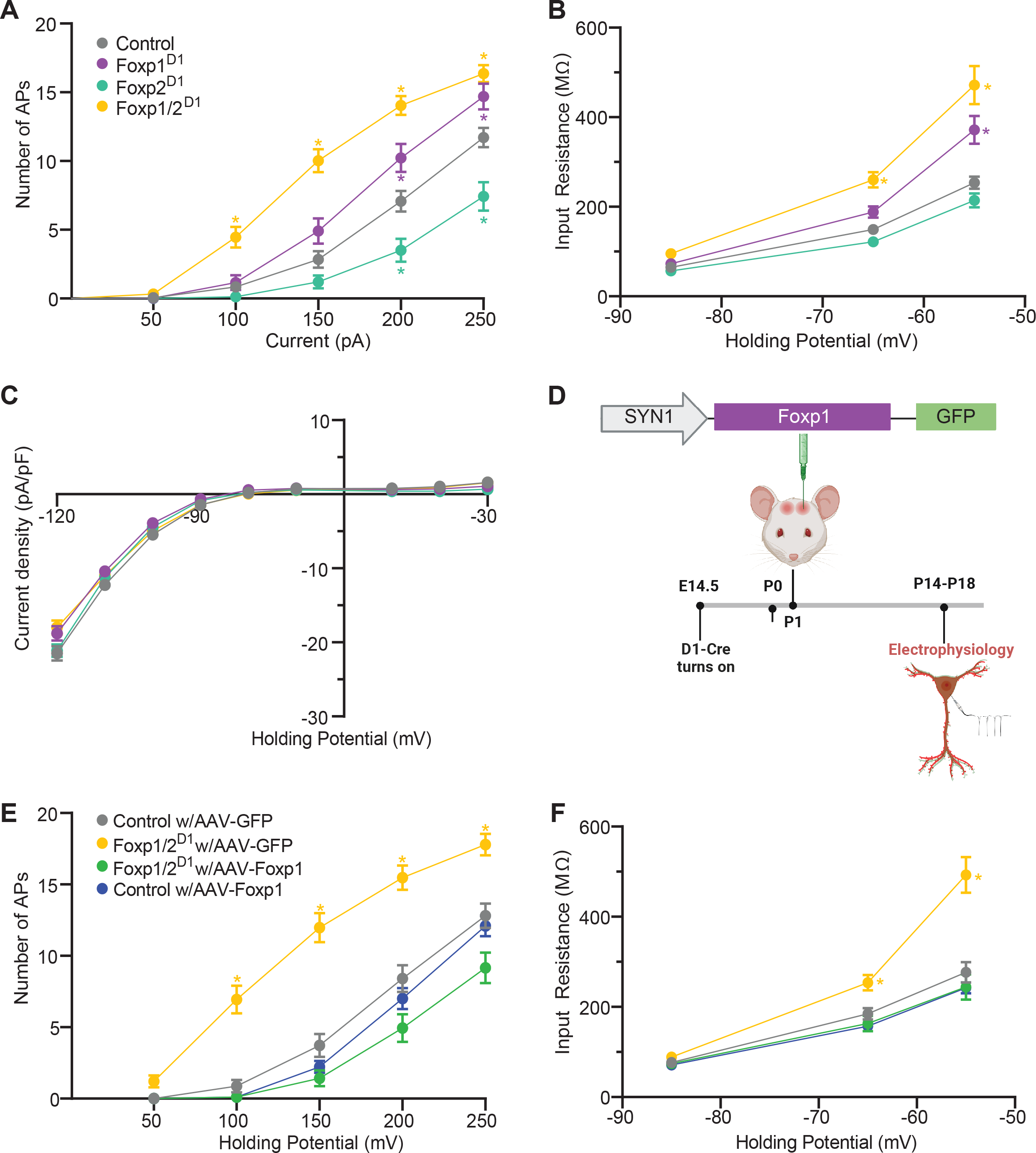
Loss of *Foxp1* results in KLeak mediated hyperexcitability with amplification by further loss of *Foxp2*. **(A)** Number of action potentials recorded under current clamp conditions. **(B)** Input resistance recorded from the same cells in the same conditions. **(C)** Contribution of KLeak channels was determined by finding the difference in current density plots. Values recorded in presence of cesium (see Figure S3D) were subtracted from those recorded in its absence (see Figure S3C) with no significant differences observed. **(D)** Schematic showing pAAV-hSYN-Foxp1-T2A-eGFP construct which was injected into mice at postnatal day 1. Dual presence of td-Tomato and GFP was used to identify which neurons had taken up FOXP1 construct. **(E)** Current clamp recordings done in mice that were injected with the FOXP1 or control construct to record number of action potentials and **(F)** input resistance. Repeated measures Two-Way ANOVA with Holm-Sidak’s post-hoc test; only significant differences between knockouts and controls are shown. *p<0.05 **(A & B)** n=56 (control), 32 (Foxp1^D1^), 40 (Foxp2^D1^), and 41 (Foxp1/2^D1^). **(C)** n=61, 43, 38, and 41. **(E & F)** n=44 (control with control virus), 34 (Foxp1/2^D1^ with control virus), 32 (Foxp1/2^D1^ with FOXP1 construct), and 19 (control with FOXP1 construct).

**Figure 7:**
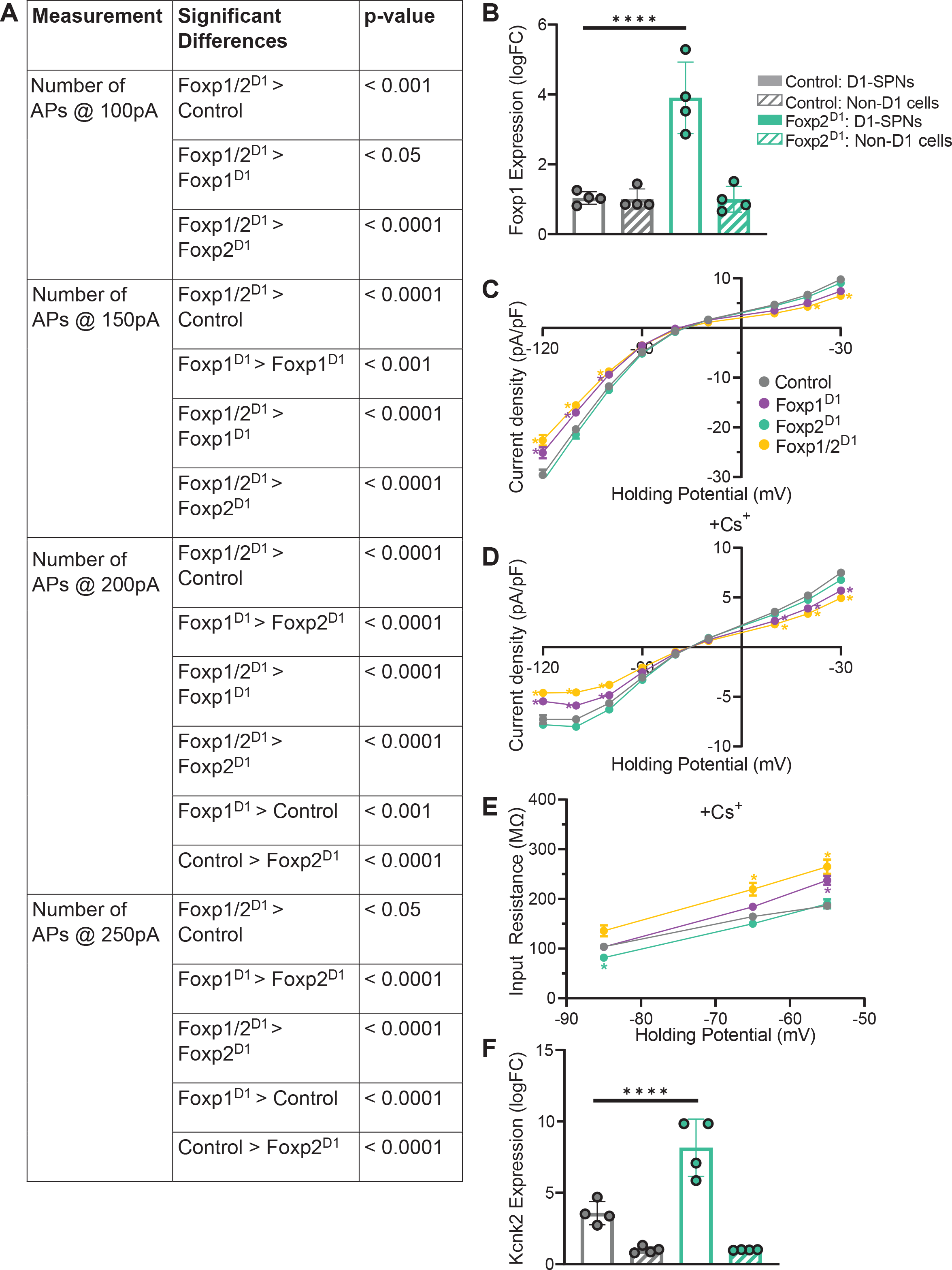
*Foxp1* and *Foxp2* mediate hyperexcitability in D1-SPNs. **(A**) Table with significant differences when comparing D1-SPN excitability between all genotypes (related to Figure 4A). **(B)** D1-SPNs from Foxp2^D1^ have greater *Foxp1* expression than those from control mice as assessed by RT-qPCR. **(C)** Current voltage (IV) plots were generated to identify KIR step density in absence and presence **(D)** of cesium to block KIR currents; both Foxp1^D1^ and Foxp1/2^D1^ D1-SPNs show differences compared to controls in both conditions. **(E)** Input resistance is increased in Foxp1^D1^ and Foxp1/2^D1^ D1-SPNs even in presence of cesium and is decreased in Foxp2^D1^. **(F)** Expression of *Kcnk2* is increased specifically in D1-SPNs from Foxp2^D1^ mice. **(B, F)** One-way ANOVA with Tukey’s post-hoc analysis, **** p < 0.0001, n=3 for all conditions. **(C-E**) Repeated measures Two-Way ANOVA with Holm-Sidak’s post-hoc test, *p<0.05, n=61, 43, 38, and 41 respectively.

### Changes in knockout mice are driven by impaired KLeak channel function

We previously reported that the hyperexcitability of D2-SPNs with loss of *Foxp1* was due to altered KIR and KLeak currents.^19^ To test the role of *Foxp1* in regulation of these currents in the D1-SPNs, we performed a separate set of experiments, this time in the presence of Tetrodotoxin (TTX) to block action potentials. For all four genotypes, we utilized a multi-step voltage protocol and measured induced currents. This was done first in control ACSF and then with subsequent wash-in of Cs^+^ to block KIR currents.^72^ Average current density versus voltage plots (IV-plots) were generated for both conditions (Figures 7C and 7D). In the absence of Cs^+^, at both depolarized and hyperpolarized potentials, Foxp1^D1^ and Foxp1/2^D1^ D1- SPNs have a current density lower than that of the controls or Foxp2^D1^ mice (Figure 7C). These differences are still detected when Cs^+^ is applied to the bath (Figure 7D). Consistent with these results, D1-SPNs in Foxp1^D1^ and Foxp1/2^D1^ mice show increased input resistance in Cs^+^ wash-in conditions (Figure S3E). We subtracted traces collected in the presence of Cs^+^ (Figure 7D) from those collected before Cs^+^ wash-in (Figure S3C) to obtain currents that are blocked by Cs^+^ (Figure 6C). We report that there are no detectable differences between controls and any of the cKOs (Figure 6C). This strongly suggests that KIR channels are unchanged upon the loss of *Foxp1* or *Foxp2* and therefore dysregulation of these channels is not driving the difference seen between genotypes.

In the presence of Cs^+^, differences in the IV plots for both Foxp1^D1^ and Foxp1/2^D1^ D1-SPNs persist (in comparison to controls) at hyperpolarized and depolarized potentials. This suggests that a voltage-insensitive current accounts for these differences. Consistent with these results, increased input resistance was observed in the D1-SPNs of these knockouts under Cs^+^ conditions (Figure 7E). We speculate a loss of voltage independent KLeak currents may account for these results, similar to what we found upon loss of *Foxp1* from D2-SPNs.^19^ Accordingly, regions associated with *Kcnk2* in the Foxp1/2^D1^ D1-SPNs were in a more repressive chromatin state and dual loss of *Foxp1* and *Foxp2* from the juvenile D1-SPNs results in downregulation of this gene (adjusted p value = 7.7 X 10^-86^). Since loss of *Kcnk2* is a factor in driving the hyperexcitability phenotype, we hypothesized that increased expression of this gene might underlie hypoexcitability of the D1-SPNs. Indeed, we observed increased *Kcnk2* in the D1-SPNs of Foxp2^D1^ in comparison to controls by both RT-qPCR (Figure 7F) and in the snRNA-seq data (adjusted p value = 1.63 X 10^-40^), suggesting that *Kcnk2* has compensatory positive regulation by Foxp1 with loss of Foxp2 in D1- SPNs. Together, our data implicate KLeak impairment downstream of loss of Foxp1 and/or Foxp2 in D1- SPNs.

### *Foxp1* is sufficient to restore D1-SPN excitability to baseline

Our findings indicate a specific role for *Foxp1* in maintenance of D1-SPN excitability. Loss of *Foxp1* resulted in hyperexcitability while upregulation of *Foxp1* in the Foxp2^D1^ D1-SPNs likely contributed to hypoexcitability. With this in mind, we considered that *Foxp1* alone may be sufficient to maintain neuronal excitability even in the D1-SPNs of double knockouts. To test this, we used an AAV-construct to reintroduce Foxp1, under the neuronal *Syn1* promoter. This construct also results in expression of green fluorescent protein for visualization purposes (pAAV-hSYN-Foxp1-T2A-eGFP). Pups were injected at P1 with this construct or a control vector also expressing GFP (AAV9-hSyn-eGFP) into the striatum (Figure 6D). As before, current clamp experiments were performed on juvenile mice between P14-P18, this time with recordings done on cells that expressed both td-Tomato and GFP. In mice with the Foxp1 construct, we no longer saw neuronal hyperexcitability of the D1-SPNs (Figure 6E). Likewise, the input resistance returned to baseline (Figure 6F). Thus, re-introduction of *Foxp1* at an early postnatal time point in Foxp1/2^D1^ mice is sufficient to rescue the hyperexcitability phenotype seen in these mice.

In summary, these results show that loss of *Foxp1* from the D1-SPNs results in hyperexcitability. The further loss of *Foxp2* from these neurons amplifies this result. We speculate that hyperexcitability may arise from loss of KLeak channels. Conversely, there is neuronal hypoexcitability observed in the Foxp2^D1^mice, seemingly driven by an increase in *Foxp1*. While both *Foxp1* and *Foxp2* play roles in maintaining D1- SPN excitability, *Foxp1* has a greater role, as evidenced by its sufficiency to rescue hyperexcitability phenotypes in Foxp1/2^D1^ mice.

### Persistent cell-type genomic changes in the striatum with loss of *Foxp1* and/or *Foxp2*

We next asked whether developmental loss of Foxp1 and/or Foxp2 results in sustained gene expression changes in D1-SPNs. Thus, we carried out snRNA-seq in striatal tissue from mice at P56. Using similar quality control filtering as before, we profiled 202,611 nuclei across all four genotypes with 3 mice per genotype (Figure 8A and 8B). We identified 11 major cell-types which were represented equally across genotypes (Figures 8C-8E). As before, to determine D1-SPN relevant changes, SPNs were subset and re-clustered for further analysis (Figure 9A). A total of 14,465 D1-SPNs were maintained in this subset and representation of each genotype within the subset SPNs remained roughly equal (Figure 8A and 8F). These subset clusters were also filtered for quality using the same standards as before (Figure 8G).

**Figure 8:**
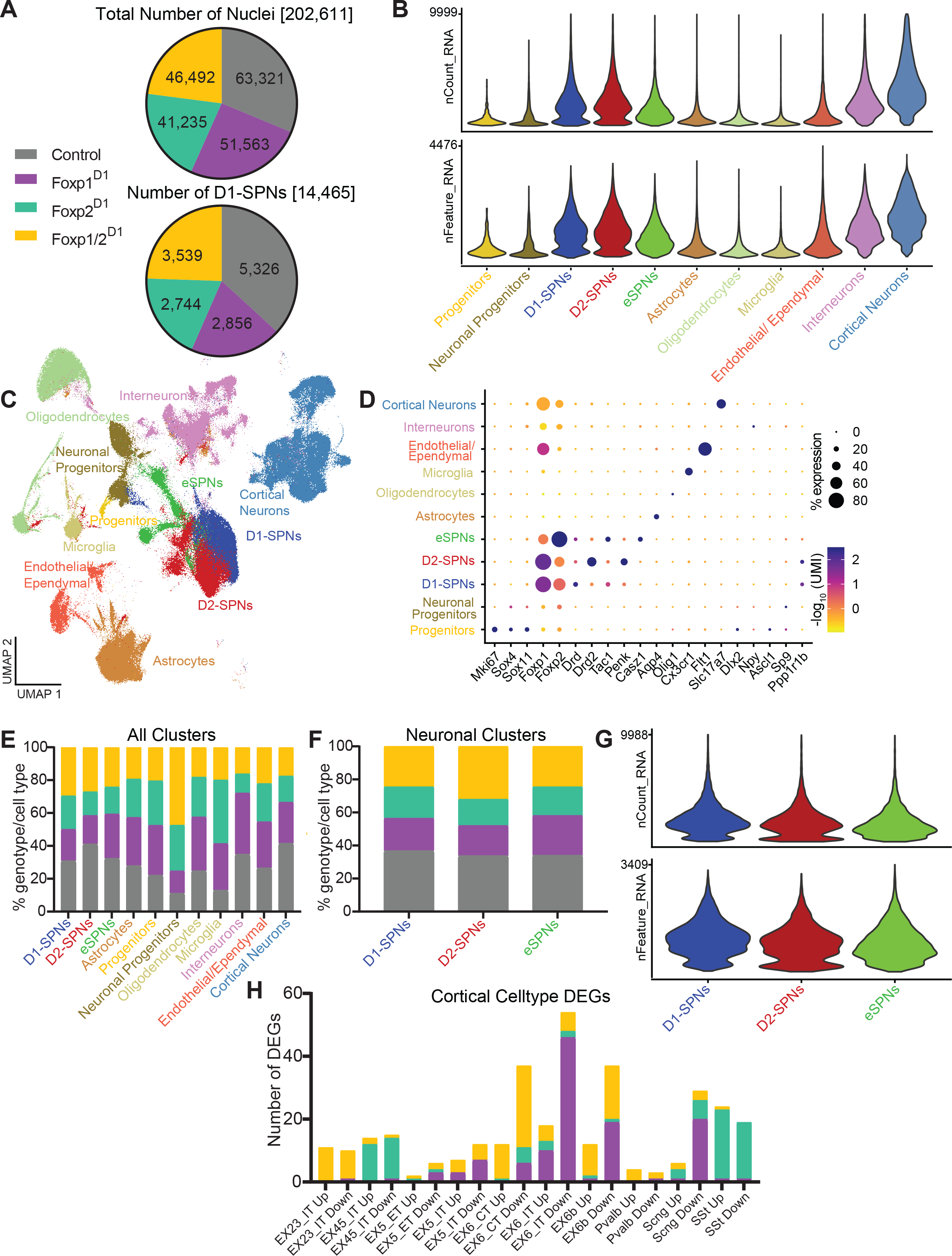
Quality control of nuclei sequenced from adult snRNA-Seq. **(A**) Pie charts showing the total number of nuclei sequenced from each genotype as well as the number of D1-SPNs profiled for each condition. **(B)** Violin plots showing the number of genes and UMI for each cell-type annotated within the dataset. **(C)** UMAP showing all of the annotated clusters from the dataset. A total of 11 major cell-types were identified. SPNs and progenitors were subsetted for further analysis. **(D)** Bubble plot showing overlap with marker genes used to annotate clusters. **(E)** Stacked bar plot showing the proportion of each genotype in all of the cell-types as well as within the neuron-only subset **(F)** that was later generated. **(G)** Violin plots showing the number of genes and UMI for the three SPN sub-types in the subset dataset. **(H)** Bar plot showing the number of DEGs in cortical cell-types in each knockout condition. EX23_IT: Excitatory cells in Layer 2/3 with intratelencephalic projections, EX45_IT: Excitatory cells in Layer 4/5 with intratelencephalic projections, EX5_ET: Excitatory cells in Layer 5 with extratelencephalic projections, EX5_IT: Excitatory cells in Layer 5 with intratelencephalic projections, EX6_CT: Excitatory cells in Layer 6 with cortico-thalamic projections, EX6_IT: Excitatory cells in Layer 6 with intratelencephalic projections, EX6B: Excitatory cells in Layer 6B, IN_PVALB: Pvalb+ inhibitory cells, IN_SCNG: Scng+ inhibitory cells, and IN_SST: Sst+ inhibitory cells.

**Figure 9:**
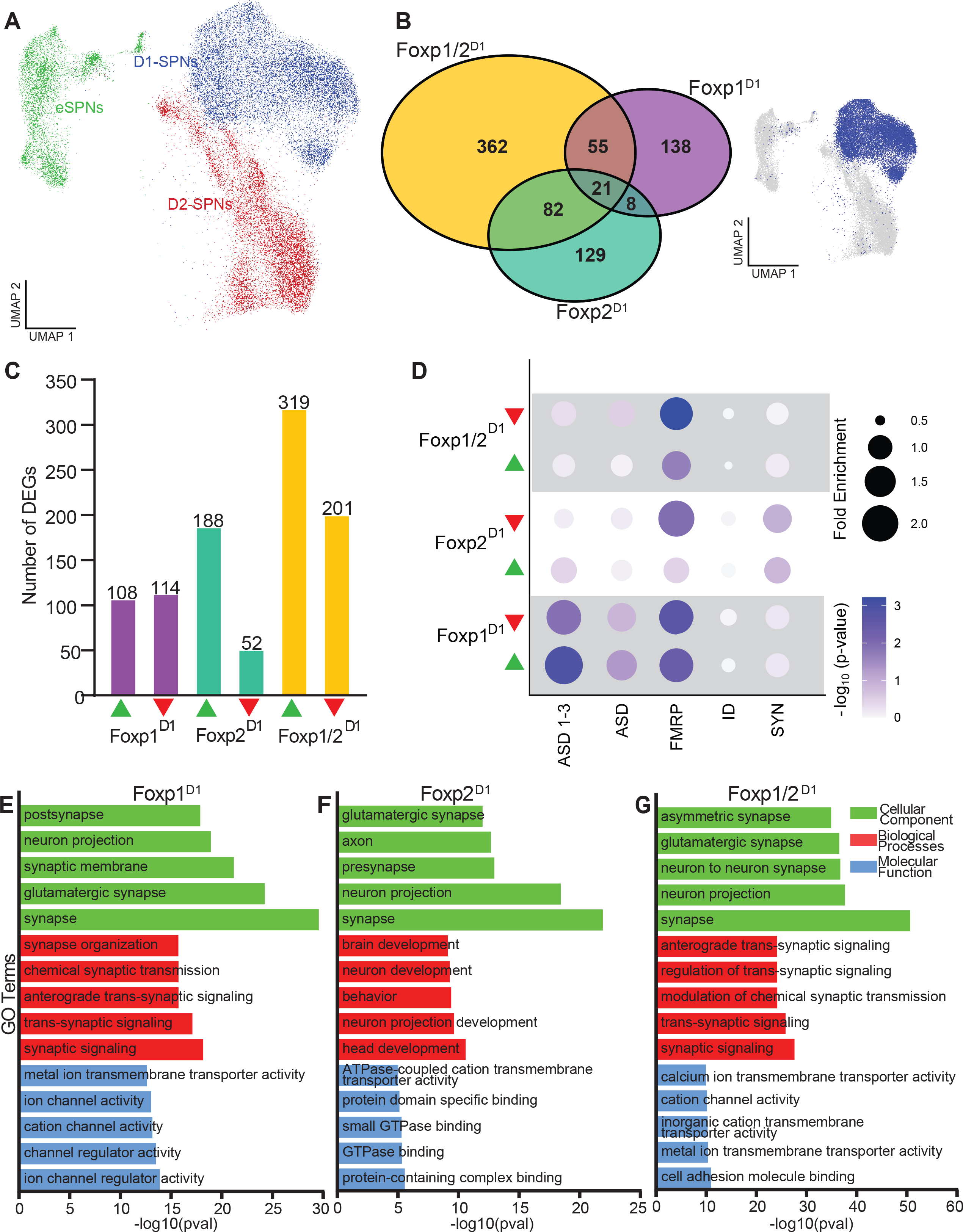
Differentially expressed genes in adults SPNs have similar biological functions as those observed in juveniles. **(A)** UMAP plot generated from the neuron-only subset with colors indicating the different annotated cell-types. D1-SPNs were used for further DEG analysis. Genes were determined to be differentially expressed from controls if they had an adjusted p-value < 0.05 and absolute logFC >|0.25|. **(B)** Semi-scaled Venn diagram showing number of unique and overlapping DEGs in each knockout condition in the D1-SPNs. **(C)** Bar plots showing the number of up- and downregulated genes in each knockout condition. **(D)** Bubble chart showing enrichment of DEGs from each knockout condition. The - log10(p-value) for each enrichment is also indicated. ASD, SFARI ASD risk genes; ASD 1-3, SFARI ASD risk genes with scores of 1-3; FMRP, Fragile X Syndrome; ID, Intellectual disability; SYN, synaptic genes. Gene Ontology (GO) analysis of **(E)** Foxp1^D1^, **(F)** Foxp2^D1^, and **(G)** Foxp1/2^D1^ DEGs reveals enrichment for terms associated with electrophysiological properties and synaptic properties.

We again determined DEGs by comparing knockouts and controls within each cell-type using the same criteria. In the D1-SPNs, we found 222 genes changing in the Foxp1^D1^ (108 up, 114 down) 240 in the Foxp2^D1^ (188 up, 52 down), and 520 in Foxp1/2^D1^ (319 up, 201 down; Figures 9B and 9C). There were 362 genes unique to the double knockout condition (70% of all Foxp1/2^D1^ D1-SPN DEGs). We hypothesize that these are genes that are normally compensated by either *Foxp1* or *Foxp2*. The magnitude of compensation in the adult tissue is much greater than what we observed in juveniles, as the number of DEGs in the Foxp1/2^D1^ is considerably greater. Again, over 60% of these genes are differentially upregulated upon the loss of both transcription factors, implicating a primarily repressive role under normal conditions.

While there was greater overlap of DEGs between conditions than was seen in juveniles, overlaps were still limited. There are 29 shared genes between Foxp1^D1^ and Foxp2^D1^ of which 23 are shared in the same direction, indicating similar types of regulation. There are 21 genes that are shared among all 3 conditions (Figure 9B and Table S1). We overlapped all DEGs with ASD-relevant gene lists and once again found enrichment for ASD risk genes (Figure 9D and Table S1). GO analysis of the DEGs for all three conditions resulted in similar terms to what we observed in the juvenile (Figures 9E-9G and Table S1). This indicates that although few specific DEGs are shared between the two time points, the loss of *Foxp1* and/or *Foxp2* affects similar gene pathways in the adult and juvenile striatum. Findings in adult snRNA-seq match what was observed in the juvenile D1-SPNs, lending further evidence to the claim that there is compensation occurring between *Foxp1* and *Foxp2*.

We also collected and annotated cortical cells in the adult dataset. Since *Foxp1*, *Foxp2*, and *Drd1* are all expressed in the cortex, we could examine whether D1-Cre affects *Foxp1* and/or *Foxp2* transcriptional programs in D1-positive cells of the cortex. We identified DEGs in 10 cortical cell-types that should express *Drd1*. Of the 30 (3 genotypes, 10 cell-types for each) comparisons that we made, 22 of the comparisons had fewer than 20 DEGs and only excitatory intra-telencephalic layer 6 neurons in the Foxp1^D1^ mice exhibited more than 50 DEGs (Figure 7H and Table S1). This indicates limited transcriptional changes in the cortex upon the loss of *Foxp1* and/or *Foxp2* using a D1-Cre.

### Loss of both *Foxp1* and *Foxp2* results in impaired motor and social behavior in mice

We next sought to determine the consequences of the transcriptional changes in the adult mice by assessing behavior in 8- to 12-week-old mice. Previous studies identified motor learning, activity, and social behavior impairments in mice with loss of either *Foxp1* or *Foxp2*.^18, 21, 34, 50, 66, 73–78^ We assessed motor learning via the accelerated rotarod. While individual deletion of one gene did not result in significant changes, the Foxp1/2^D1^ mice showed a significant impairment compared to controls (Figure 10A). There were no differences in grip strength between any of the genotypes (Figures 11A and 11B). Like motor learning, motor activity in the open field assay was significantly impaired with loss of both *Foxp1* and *Foxp2* (Figure 10B). In line with previous reports, there was no anxiety phenotype as assessed by the percentage of time spent in periphery of the open field box (Figure 11C).^50, 77^ Together, these findings are consistent with the hypothesis that *Foxp1* and *Foxp2* can compensate for each other to regulate motor activity. Findings with deletion of only *Foxp1* also match our previous findings.^18^

**Figure 10:**
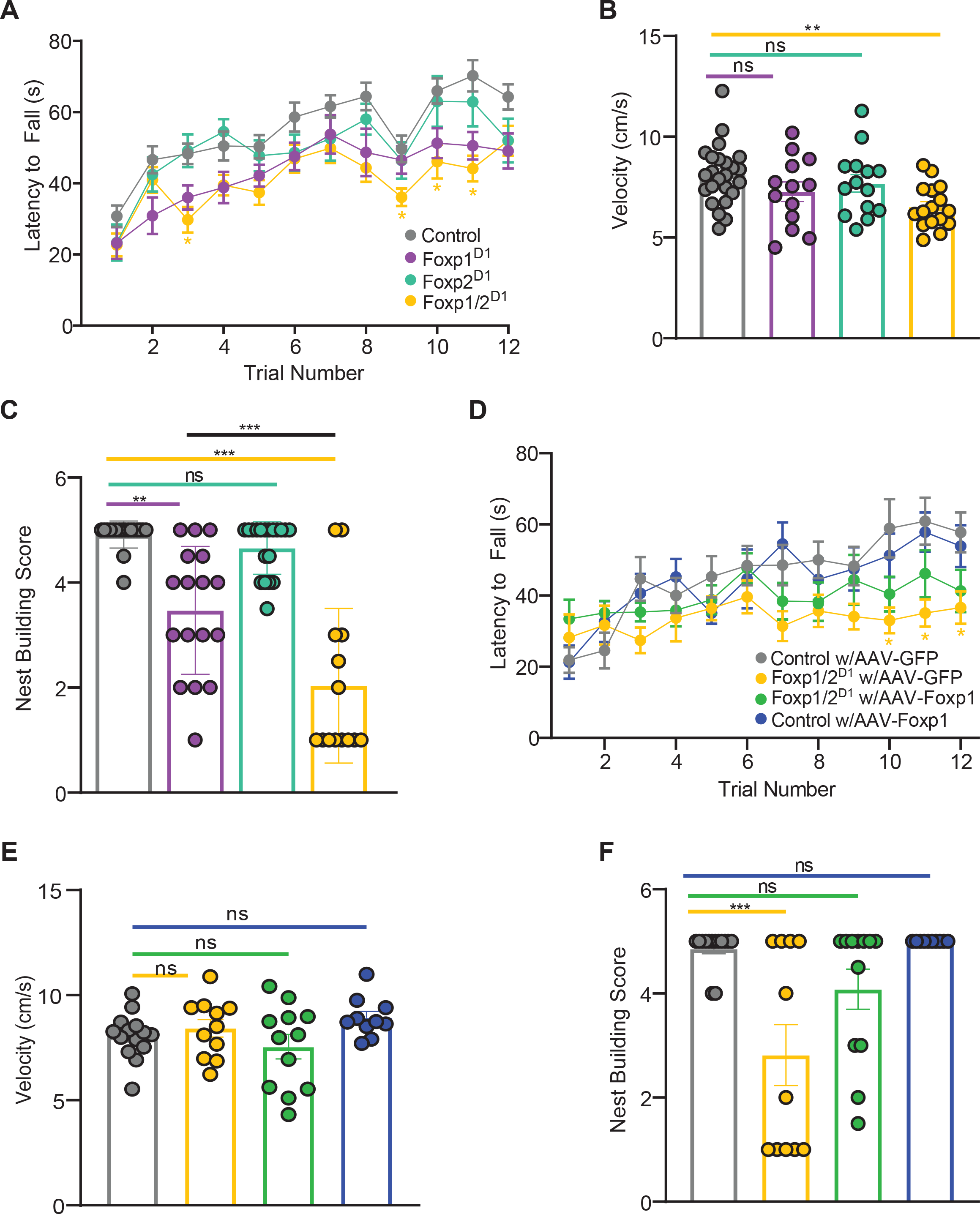
Loss of *Foxp1* and *Foxp2* results in impaired motor and social behavior. **(A**) Foxp1/2^D1^ mice show motor learning deficit as assessed by latency to fall using the rotarod paradigm and in **(B)** motor activity measured as observed by amount of movement during 5-minute open field assay. **(C)** Nest building quality was assessed after single housing for 24 hours where Foxp1^D1^ had an impairment. Impairment amplified in Foxp1/2^D1^. AAV-mediated re-expression of *Foxp1* restored measures to baseline in rotarod **(D)**, motor activity **(E)**, and nest building **(F)**. **(A-C)** Behavior performed in control, Foxp1^D1^, Foxp2^D1^, and Foxp1/2^D1^. **(D-F)** Performed in control or Foxp1/2^D1^ mice with either control AAV9-hSYN1-eGFP or pAAV-hSYN-Foxp1-T2A-eGFP construct. Two- (**A&D)** or One- **(B, C, E, & F)** Way ANOVA with Tukey’s post-hoc analysis used to determine significance. *p<0.05, **p<0.01, ***p<0.001. n=25 (control), 13 (Foxp1^D1^), 14 (Foxp2^D1^), and 16 (Foxp1/2^D1^) for **(A&B)**, n=15, 13, 18, and 12 respectively for **(C)**, and n=14, 11, 12, and 10 respectively for **(D-F)**.

**Figure 11:**
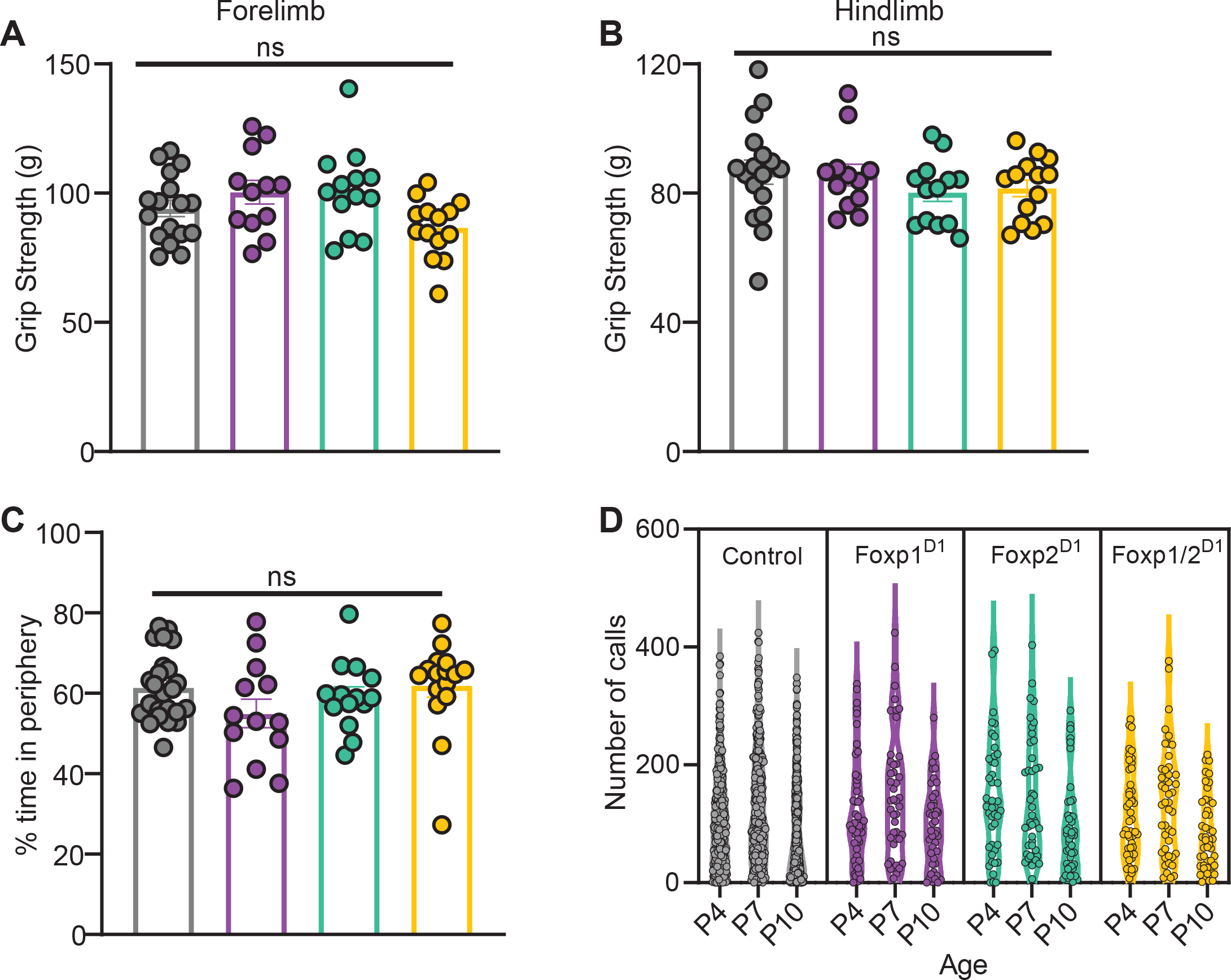
Loss of *Foxp1* and/or *Foxp2* has no effect on limb strength or anxiety. No deficits were seen in either fore- (**A**) or hind-(**B**) limb grip strength. **(C)** No changes were seen in anxiety levels as assessed by time spent in periphery during the 5-minute open-field assay. **(D)** Violin plots showing the number of pup isolation ultrasonic vocalizations made by mice of all four genotypes at postnatal days 4, 7, and 10. No changes were observed when comparing number of calls made at each time point across genotypes. One- (**A-C**) or Two- (**D**) Way ANOVA with Tukey’s post-hoc analysis used to determine significance. n=17, 12, 13, and 14 respectively for (**A-C**), and n=310, 46, 44, and 47 for (**D**).

To assess social behavior in adult mice, we tested nest building, a communal behavior that can be reproducibly quantified in rodents.^18, 79^ While the Foxp2^D1^ mice have no deficit in nest building, Foxp1^D1^ mice created significantly less well-developed nests, in line with our previous findings.^18^ The Foxp1/2^D1^ mice, however, demonstrated even significantly greater nest building impairments (Figure 10C). This indicates that *Foxp2* is not sufficient to fully rescue the effects of *Foxp1* deletion in this social behavior. Instead, the loss of *Foxp2* in addition to *Foxp1* further exacerbates the nest building deficits. We also examined pup isolation ultrasonic vocalizations (USVs), which are calls made by mouse pups to elicit maternal care. While we previously reported that Foxp1^D1^ mice made fewer calls than controls^18^, we were unable to replicate this finding in the same genotype in this study. Furthermore, neither the Foxp2^D1^ nor the Foxp1/2^D1^ mice showed any deficiencies in the number of USV calls (Figure 11D). These findings indicate that *Foxp1* and *Foxp2* have mostly redundant functional roles in regulating motor and social behaviors in adult mice. However, *Foxp1* has a greater impact compared to *Foxp2*, as indicated by the nest building impairment observed in Foxp1^D1^ mice.

### Viral mediated re-expression of *Foxp1* is sufficient to restore behavioral impairments

We next asked whether expression of *Foxp1* is sufficient to restore behavioral deficits in Foxp1/2^D1^ mice, similar to its restorative effects on neuronal excitability. To this end, we injected either a control or Foxp1 expressing construct (as previously described) bilaterally into the striatum of pups at P1 and assessed behavior when mice were 8-12 weeks old. The double knockouts with restored Foxp1 no longer show an impairment in rotarod (Figure 10D), motor activity (Figure 10E), or nest building (Figure 10F). Thus, re- introduction of *Foxp1* at an early postnatal time point was sufficient to restore behavioral impairments typically observed in adult Foxp1/2^D1^ mice. It should be noted, however, that the Foxp1/2^D1^ mice injected with the control virus no longer show a deficit in the open field either.

## Discussion

In this study, we examined overlapping and unique functions of the transcription factors Foxp1 and Foxp2 in striatal D1-SPNs. Using conditional mouse models to specifically knockout *Foxp1*, *Foxp2*, or both from the D1-SPNs, we found that the combined loss of both genes amplified impairments in KLeak mediated hyperexcitability as well as motor and social behaviors, indicating compensatory roles of these two transcription factors. Our data from single nuclei transcriptomics supports this conclusion with the observation that forkhead motifs are enriched in regions of chromatin that are more accessible in D1-SPNs in double cKO mice. We propose two primary mechanisms of compensation. In the first scenario, either *Foxp1* or *Foxp2* can regulate a given gene. The loss of one transcription factor results in maintained regulation by the other and only upon loss of both factors is gene expression is altered (represented by DEGs unique to Foxp1/2^D1^). In the second scenario, loss of one transcription factor leads to increased regulation of the target gene by the remaining factor, but decreased regulation upon loss of both (represented by DEGs overlapping in the same direction between single cKOs and moving in the other direction in the double knockouts). Based on the limited overlap of transcriptional targets between the single knockouts, we suggest the first mechanism as the primary method through which compensation occurs (Figure 12).

**Figure 12:**
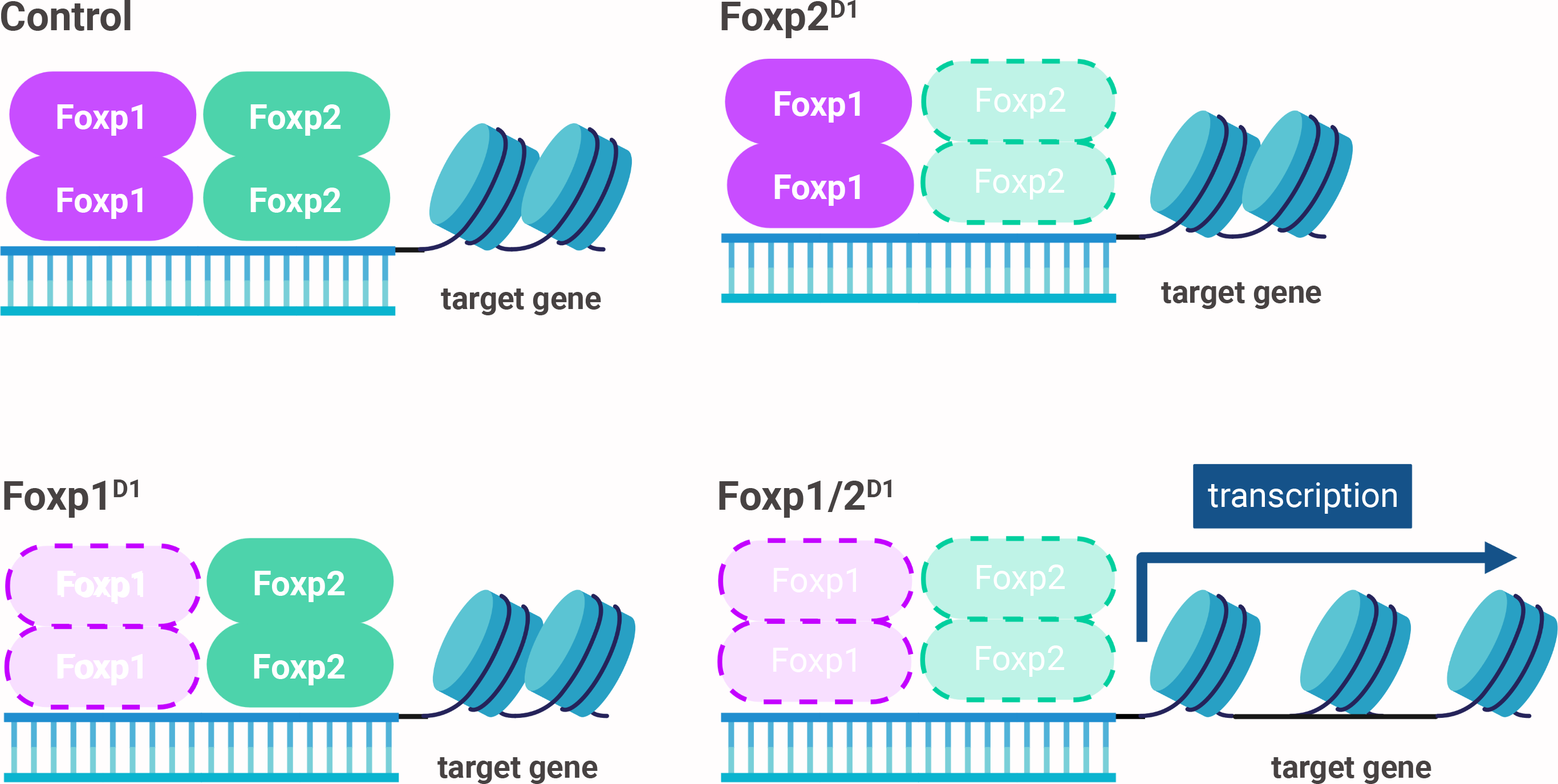
*Foxp1* and *Foxp2* largely compensate to maintain repression of target genes. We propose one mechanism of compensation wherein both *Foxp1* and *Foxp2* bind to and regulate similar targets. Upon loss of one transcription factor, the remaining transcription factor takes over regulation of these shared targets and maintains chromatin in a closed state. Upon loss of both factors, we observe dysregulation of target genes and altered chromatin state.

While our results provide evidence of compensation, there are still some possibilities about compensation that will require further work to answer. One of the transcription factors could acquire targets that are normally maintained by its paralogous gene. Or upon the loss of one transcription factors, the other could increase its regulation without changing binding. A further possibility is that target genes are preferentially bound by a heterodimer of Foxp1 and Foxp2, but regulation is maintained by a homodimer upon loss of one. The similarity in DNA binding motifs of the two genes makes it challenging to definitively distinguish between these possibilities. Another method of compensation may occur through co-regulation of other transcription factors. For example, it is only in the Foxp1/2^D1^ D1-SPNs that we observed an enrichment of Mef2 motifs within DARs. The Mef2 family of genes, which are known to be downregulated by *Foxp1* and *Foxp2*,^21, 36, 51, 71^ may thus be regulated in a compensatory manner. Upon loss of both Foxp genes, we propose that there is dysregulation of not only Foxp1/Foxp2 targets, but also Mef2 family transcriptional targets. *Mef2a, Mef2c,* and *Mef2d* are all upregulated in juvenile (adjusted p value = 4.23 X 10^-54^, 5.04 X 10^-32^, and 1.81 X 10^-29^ respectively) while only *Mef2c* was upregulated in the adult (adjusted p value =.0008) Foxp1/2^D1^ D1-SPNs, adding further evidence to this possibility.

While there is likely large-scale compensation occurring, we also identified genes that are uniquely regulated by either Foxp1 or Foxp2 where there is no robustness between the two transcription factors. The non-overlapping DEGs in the single knockouts represent genes that are only regulated by either *Foxp1* or *Foxp2*. This indicates that despite their similarities, there are distinct targets for each transcription factor. Evidence of this is seen in the Foxp1^D1^ mice that have electrophysiological and behavioral phenotypes not seen in controls, indicating that *Foxp2* is not fully compensating for *Foxp1*.

D1-SPNs in Foxp1^D1^ mice have KLeak mediated increases in neuronal excitability, which is amplified in the Foxp1/2^D1^ mice. Thus, while loss of *Foxp1* is sufficient to develop a phenotype, *Foxp2* still plays a role as evidenced by the exacerbated hyperexcitability. The hypoexcitability observed in Foxp2^D1^ D1-SPNs represents a scenario wherein *Foxp1* “over” compensates and increases regulation of genes involved in mediating neuronal excitability. In support of this idea, we found that *Kcnk2*, a gene we have previously identified as playing a role in mediating hyperexcitability in D2-SPNs^19^, was specifically upregulated in the Foxp2^D1^ D1-SPNs. Furthermore, an AAV-mediated over-expression of Foxp1 is sufficient to restore neuronal excitability in Foxp1/2^D1^ mice, indicating that *Foxp1* is sufficient to compensate for loss of *Foxp2*. Our study focused on the potential contribution of KIR and KLeak channels to neuronal hyperexcitability. However, there are other types of potassium channels that show differential expression upon loss of both *Foxp1* and *Foxp2*, including the potassium voltage-gated channels *Kcnip1* and *Kcnb1*. Interestingly, variants in the latter have been implicated in apraxia of speech, a phenotype also seen in human patients with mutations in *Foxp1* or *Foxp2*.^57, 58, 80^ Further studies can be done to determine how the downregulation of these genes may impact the phenotypes we observe.

We report that motor learning and motor activity were only impaired upon loss of both genes indicating that *Foxp1* and *Foxp2* can robustly compensate to maintain proper function of the circuits dictating motor behavior. D1-SPNs are generally involved in activating movement and here we report that loss of *Foxp1* and *Foxp2* inhibited movement in the open field. Thus, each gene can maintain proper SPN function for overall movement with impairments only arising when both genes are deleted. The same effect was observed when we investigated motor learning. Foxp1 on its own was sufficient to restore motor learning to baseline levels in the Foxp1/2^D1^. Unlike motor behavior, social behavior (as measured by nest building) is impaired upon the loss of *Foxp1* alone, highlighting a requirement for *Foxp1* in that circuit. However, the impairment is exacerbated upon further loss of *Foxp2*, indicating that *Foxp2* synergizes with *Foxp1* function in this behavior.

Using single-nuclei transcriptomics, we identified hundreds of genes that are regulated by both *Foxp1* and *Foxp2*. Given the important role of these transcription factors in human brain disorders, it will be important to determine the contribution of subsets of differentially expressed genes to disease-relevant D1- SPN development and function. For example, members of the neurexin family of genes (*Nrxn1, Nrxn2,* and *Nrxn3*), which have been implicated in ASD, were differentially regulated.^81, 82^ Recent studies report that loss of *Nrxn1* impaired synaptic connections onto the SPNs^83, 84^ but their roles in D1-SPN development are yet to be studied. We also identified many genes crucial for regulating the G-protein signaling cascade which is required for proper D1-SPN function; these include *Pde1b, Pde1c, Prkca* (protein kinase C alpha), and *Prkcb* (protein kinase C beta).^85–92^ Adenylyl cyclase 5 (*Adcy5*), which also mediates GPCR signaling and has previously been found to affect rotarod performance^93^, is downregulated in adult Foxp1/2^D1^ mice. Another recent study identified upregulation of *Dyanactin1* (*Dctn1*), which encodes a subunit of dynein that is important for regulation of proper protein motor homeostasis in striatal neurons, in mice with *Foxp2* mutations.^94^ In juvenile Foxp1/2^D1^ D1-SPNs, *Dctn1* had a more permissive chromatin state and was upregulated (adjusted p value = 1.12 X 10^-12^). Thus, alterations in these genes result in striatal dysfunction and provide compelling avenues for follow up studies downstream of Foxp1 and Foxp2 in the striatum.

In summary, we find that Foxp1 and Foxp2 have compensatory roles in the D1-SPNs. This is the first study to investigate the interaction between these transcription factors in a neuronal population that expresses both genes. D1-SPNs are crucial for motor behaviors, and it is only upon the loss of both genes that mice showed impairments in motor learning and motor activity. While nest building was impaired upon the loss of *Foxp1*, the impairment was amplified upon the further loss of *Foxp2*, again indicating some level of compensation. The same trend was observed in KLeak mediated D1-SPN hyperexcitability. Reintroducing Foxp1 on its own was sufficient to restore behavioral and electrophysiological phenotypes to baseline in Foxp1/2^D1^ mice, again indicating that *Foxp1* can compensate for the loss of *Foxp2*. Data from single-nuclei transcriptomics supports the compensatory interactions of both genes as evidenced by hundreds of differentially expressed genes and differentially accessible regions in the D1-SPNs. Overall, the results presented here provide insight into how the paralogous transcription factors Foxp1 and Foxp2 can work together to mediate D1-SPN development and function.

## Acknowledgements

Our sincerest thanks to Dr. Peter Tsai, Dr. Maria Chahrour, and Dr. Todd Roberts for providing critical feedback on the manuscript. We thank Dr. Shin Yamazaki and the Neuroscience Microscopy Facility, supported by the UT Southwestern Neuroscience Department and the UTSW Peter O’Donnell, Jr. Brain Institute. We would like to thank Dr. Shari Birnbaum at the UTSW Rodent Behavior Core for help performing the open field analysis. We would also like to thank Dr. Eric Plautz, Dr. Denise Ramirez, and the UTSW Neuro-Models Facility for providing access to the equipment for rotarod and grip strength. Finally, we would like to thank Dr. Benjamin Arenkiel, Dr. Joshua Ortiz, and the Baylor College of Medicine Optogenetics and Viral Design/Expression Core for designing the AAV-SYN1-FOXP1-GFP construct. G.K. is a Jon Heighten Scholar in Autism Research and Townsend Distinguished Chair in Research on Autism Spectrum Disorders at UT Southwestern. This work was supported by grants from NIMH (MH126481, MH102603), NINDS (NS126143, NS115821), the Simons Foundation (573689, 947591), and the James S. McDonnell Foundation 21^st^ Century Science Initiative in Understanding Human Cognition – Scholar Award (220020467) to G.K. This work was further supported by the NIH/NIMH (grant T32-HL 139438 to N.I.A) and the NIH/NINDS (grant NS117030-01 to N.I.A.).

## Author Contributions

N.I.A. and G.K. designed the study and wrote the paper. N.I.A. performed single-nuclei RNA and ATAC sequencing experiments, and mouse behavior experiments. A.G.A. contributed to initial study design and aided in single-nuclei RNA sequencing experiments. J.G. and N.K. designed the electrophysiology experiments and aided in writing the paper. N.K. performed electrophysiology experiments and neonatal AAV injections. N.K. and N.I.A. performed FACS and qPCR experiments. A.K. performed pre-processing and analysis of all single-nuclei RNA and ATAC sequencing data.

## Resource availability Lead contact

Further information and requests should be directed to and will be fulfilled by the Lead Contact, Genevieve Konopka.

## Materials availability

Animals and materials generated from this study are available from the lead contact with a completed Materials Transfer Agreement.

## Data and Code Availability

All acquired data and code are available upon request to the lead contact.

## Experimental model and subject details

Experiments were performed in accordance with procedures approved by UT Southwestern’s Institutional Animal Care and Use Committee (IACUCC #2016-101825). All mice were C57Bl/6J. *Foxp1^flox/flox^, Drd1a*- *Cre* (262Gsat), and *Drd1-tdTomato* mice were used as previously described.^18^ *Foxp2^flox/flox^*mice were obtained from Jackson Laboratories (Strain# 026259).^95^ *Foxp1 ^flox/flox^* and *Foxp2 ^flox/flox^* mice bred to the *Cre* and reporter lines were maintained separately for the generation of single-knockout mice while a separate line was used to generate mice that were homozygous for both *Foxp1 ^flox/flox^* and *Foxp2 ^flox/flox^*. Both male and female mice were used in all experiments in equal numbers. All mice were located in the same room in the mouse facility and were maintained on a 12-hour light on/off schedule.

## Method details

### Data analysis for behavior and electrophysiology

For behavior data, sample number is the number of animals. An equal number of male and female mice were used for experiments. For electrophysiology, sample number is the number of neurons. All data are plotted as mean ± standard error. Unless otherwise indicated, a one- or two-way analysis of variance (ANOVA) with Tukey’s or Holm-Sidak’s post-hoc multiple comparisons test was used.

### Behavior

#### Nest Building

Nests were scored as we previously published.^18^ After weaning, mice were co-housed with other mice of the same sex. Once they reached >6 weeks of age, the mice were then separated and singly housed in new cages that had an intact nestlet material. Mice were then kept in the cage overnight and after approximately 18 hours, the nests were scored to assess quality on a scale from 1-5 as was previously described.^79^

#### Open Field

Mice were individually placed in a 16’’ X 16’’ Plexiglass box and allowed to explore the arena for 10 minutes. Videos of each mouse were scored for total distance moved and amount of time spent in periphery using the EthoVision XT software package (Noldus). Tests were performed by the UTSW Rodent Behavior Core directed by Dr. Shari Birnbaum. Average velocity was determined by dividing the total distance moved by the amount of time spent moving.

#### Rotarod

Following previously published methods, adult mice were brought to the room with the testing apparatus, weighed, and given 30 minutes to habituate.^18, 21^ They were then placed on a textured rod within individual lanes of a Series 8 IITC Life Science rotarod. The rod was programmed to accelerate from 4 to 40 rpm within a 5-minute time frame. Each mouse was placed, facing forward, on the rod before starting the test. Sensors located below the rod were activated when the mouse fell off the rod. If the mouse made one full rotation holding onto the rod, the sensor was manually activated, and the mouse was taken off the rod. Once all mice in a test had completed the paradigm (maximum of 5 at a time), latency to fall and maximum revolutions per minute at fall were recorded. Mice were then placed back in their home cage, rods and sensors were cleaned with 70% ethanol, and then the next set of mice were tested. Mice of opposite sexes were not tested at the same time. Mice were tested for three consecutive days with four trials per day, separated by 10-minute intervals.

#### Grip Strength

Following previously published protocols, forelimb and hindlimb grip strength were measured at least one day after completion of the rotarod.^18, 21^ The Chatillon Force Measurement equipment was used to record the amount of strength it took to pull the mouse off a mesh wire. Fore (or hind) limbs of each mouse were placed onto a mesh wire meter and then pulled away using constant force. Five consecutive measurements were recorded for both forelimbs and hindlimbs. An average grip strength for both forelimbs and hindlimbs was obtained per mouse.

#### Neonatal ultrasonic vocalizations

USVs were recorded as previously described.^18, 21, 73^ Briefly, pups were isolated form dams at P4, P7, and P10 and placed into a soundproof container that was equipped with an UltraSoundGate condenser microphone. The recordings were made using Avisoft Bioacoustic software USVs were recorded for 3 minutes and then pups were returned to their dams. Analysis of sound spectrograms was performed using an established MATLAB script.^96^

### Single-nuclei RNA-Sequencing (snRNA-Seq)

#### Tissue Collection and Processing (P56)

P56 mice were sacrificed by rapid dissection and brains were quickly removed and placed in ice-cold 1X PBS. Using a brain matrix with 1mm markings, striatal sections were obtained, as determined by visual cues indicating the presence of the striatum. Using forceps, the striatum was separated from the cortex, flash frozen, and stored in -80C. Nuclei isolation for snRNA-Seq was adapted from our own published protocols optimized for adult mouse striatum.^18, 97^ Briefly, tissue was homogenized in a glass Dounce tissue grinder (25 times with pestle A, 25 times with pestle B; Sigma, Cat#D8938) in 2mL ice-cold EZ Nuclei Lysis Buffer (Sigma, Cat #NUC-101). Samples were incubated on ice for 5 minutes with an additional 2mL ice-cold EZ lysis buffer and then centrifuged at 500 X g for 5 minutes at 4C, washed with 2mL ice-cold EZ lysis buffer, and incubated on ice for another 5 minutes before being spun again. Nuclei were then washed in 500uL Nuclei Suspension Buffer (NSB; 1X PBS, 0.01% Ultrapure BSA, and 0.1% RNase inhibitor) and mixed with 900uL nuclei sucrose cushion buffer (Sigma, Cat #NUC-201). This mixture was layered on top of another 500uL of sucrose cushion buffer and spun at 13,000 X g for 45 minutes at 4C. The supernatant was then discarded, the pellet was resuspended in 60uL NSB, and filtered through FLOWMI tip strainer (Bel-Art Products, Cat# H13680-0040); more NSB was added as needed to help samples go through the FLOWMI tip. An aliquot was stained with Trypan Blue and counted under a microscope to determine concentration (targeting 700-1,200 nuclei/uL). Samples were stored on ice until library generation. Libraries were prepared using the 10X Genomics Single Cell Reagent Kits v3 and v3.1 protocol targeting 10,000 nuclei total per sample.^98^ A total of 12 mice (3 mice/genotype) were prepped in 5 batches. Libraries were sequenced on an Illumina NovaSeq via the McDermott Sequencing Core at UT Southwestern.

#### Tissue Collection and Processing (P9)

Striatal tissue for P9 mice was collected using our previously published methods.^18^ Mice were sacrificed by rapid decapitation and brains were removed and placed in ice-cold artificial cerebrospinal fluid (ACSF; 126 mM NaCl, 20 mM NaHCO3, 20 mM D-Glucose, 3 mM KCl, 1.25 mM NaH2PO4, 2 mM CaCl2, 2 mM MgCl2) bubbled with 95% O2 and 5% CO2. 500uM coronal sections were made in ACSF using a VF- 200 Compresstome (Precisionary Instruments) and transferred to another chamber filled with ACSF. From 3 striatal sections, 6 punches of striatal tissue were collected (2 per hemisphere), flash frozen, and stored in -80C. Nuclei were then isolated in a similar fashion as adults, with only one 5-minute centrifugation in EZ Nuclei Lysis buffer. No sucrose cushion was used. Again, samples were filtered through the FLOWMI tip strainer, stained with Trypan Blue, counted under a microscope, and stored on ice until library generation. 12 mice were prepped in 5 batches. Libraries were sequenced as described above.

#### Pre-processing of sequencing data

Raw sequencing data was acquired from the McDermott Sequencing Core at UT Southwestern in the form of binary base call (BCL) files. BC files were de-multiplexed with the 10X Genomics i7 index (used during library preparation) using Illumina’s bcl2fastq v2.19.1 and the *mkfastq* command from 10X Genomics CellRanger v3.0.2 suite. Resulting FASTQ files were checked for quality using FASTQC (v0.11.5).^99^ A reference mouse genome-annotation index was built using mouse genome (GRCm38p6) and Gencode annotation (vM17) with *mkref* command from 10X Genomics CellRanger v3.0.2 suite. Extracted and quality passed Extracted FASTQ reads were then aligned to reference mouse genome-annotation index and raw count tables were generated using *count* command from 10X Genomics CellRanger v3.0.2 suite.

Cellbender was then run on the raw unfiltered count tables to discard potential ambient RNA.^100^ Potential doublets were identified with DoubletFinder using filtered count tables generated by CellBender.^101^ This produced an expression matrix containing cells as rows and genes as columns which was used for downstream analysis.

#### Clustering analysis

Cleaned count tables were used to run Seurat pipeline following vignette (https://satijalab.org/seurat/articles/pbmc3k_tutorial.html). Samples across genotypes were processed through Harmony to remove covariate effects such as batch and sex.^102^ Harmonised Seurat objects were integrated following vignette (https://satijalab.org/seurat/articles/integration_introduction.html). Integrated data was clustered to identify clusters. Cell-type annotation was performed using known marker genes and running Fisher exact test against cell-type marker genes identified in a previous dataset.^69^ Furthermore, nuclei corresponding to SPN classes were sub-clustered to resolve specific SPN sub-types (D1, D2, and eccentric SPNs) with higher granularity.

#### Differential gene expression analysis

For differentially expressed genes, nuclei corresponding to each SPN class were grouped by genotype. Genes with significant differential expression within D1-SPNs (or D2- or eccentric) of knockouts were identified (filtered using adjusted p-value < 0.05, |logFC| > 0.25) when compared to control samples using MAST-GLM.^103^ Differential expression reflected changes in gene expression in SPN population across genotypes.

#### Gene ontology

Gene ontology of for DEGs was performed using ToppGene.^104^ We used Gene Ontology an Kyto Encyclopedia of Genes and genomes databases. Expressed genes in the D1-SPNs (17,851 for juvenile and 16,542 for adult) were used as background.

#### Overlap with other gene databases

ASD-associated genes were downloaded from SFARI Gene database.^105^ ASD (1–3) are ASD genes with a score between 1 and 3. Fragile-X associated genes were downloaded from Darnell et al 2011.^106^ Intellectual disability gene dataset was downloaded from Chen et al 2018.^107^ Synaptic genes were downloaded from Pirooznia et al 2012.^108^

### Single-nuclei Assay for Transposase-Accessible Chromatin Sequencing (snATAC-Seq)

Striatal tissue from P9 mice were collected in the same manner as they were for snRNA-seq samples. Nuclei were isolated using similar methods as above but using different nuclei wash buffers specialized for ATAC-Seq (10mM Tris pH 7.4, 10mM NaCl, 3mM MgCl2, 10% BSA, and 1% Tween-20). After counting, nuclei were diluted using the 20X Nuclei Buffer (component of 10X ATAC-Seq kit) before proceeding with the 10X Genomic Single Cell ATAC Kit v1.1 and v2 protocol.^98^ Between 7,000-10,000 nuclei were targeted per sample. A total of 12 mice (3/genotype) were prepped in 3 batches. Libraries were sequenced as described above.

#### Pre-processing of sequencing data

Raw BCL data for sequenced libraries were received from McDermott Sequencing Core at UT Southwestern and FASTQ files were extracted using *cellranger-atac* from 10X Genomics CellRanger-ATAC v2.0.0 suite and bcl2fastq (v2.20.0). FASTQC (v0.11.5) was run on extracted FASTQ files to check the reads quality.^99^ Reference mouse genome-annotation index was built using mouse genome (GRCm38p6) and Gencode annotation (vM17) with *cellranger-atac mkref* from the 10X Genomics CellRanger-ATAC v2.0.0 suite. Fragment files were further used to identify potential doublets using ArchR (v1.0.2).^109^ BAM files were converted to BED files using *bedtools bamtobed* (v2.29.2).^110^ BED files were then used to call peaks using *MACS2 callpeak.*^111^ Using Signac (v1.9.0) and Seurat (v4.3.0), chromatin assays and Seurat objects were generated per biological replicate per genotype for shared peaks across all genotype replicates.^112–114^ Signac pipeline was run following vignette (https://stuartlab.org/signac/articles/mouse_brain_vignette.html) to generate gene activity matrix. Individual Seurat objects processed through Signac were then filtered to remove low quality nuclei (total fragments in peaks > 1,500; total fragments in peaks < 100,000; percent reads in peaks > 10%; nucleosome signal < 2; TSS enrichment > 2; blacklist ratio < 0.03) and potential doublets scored by ArchR.

#### Clustering analysis and identification of differentially accessible regions

Samples per genotype were first merged and then processed through Harmony (v0.1.0) to remove effect of covariates such as batch and sex.^102^ Harmonized Seurat objects per genotype were then integrated and data were clustered to identify clusters. Clusters were annotated using age-matched single-nuclei RNA sequencing data (P9) with *TransferData* function available as a part of Seurat following vignette (https://satijalab.org/seurat/articles/integration_mapping.html). Nuclei belonging to D1-SPN class were used to identify differentially accessible regions (DARs) across genotypes. Significant DARs (adjusted p- value <= 0.05, absolute log fold change >= 0.1375) were identified by comparing each knockout against control genotypes using *FindMarkers* command and likelihood ratio test (LR) following Signac vignette (https://stuartlab.org/signac/articles/mouse_brain_vignette.html). Enriched motifs were then identified for DARs following Signac’s *FindMotifs* command (https://stuartlab.org/signac/articles/motif_vignette.html). ^114^ All compared peaks across a pairwise comparison were used as background for motif analysis and motifs with fold enrichment > 1.75 were retained.

### Electrophysiological recordings

All experiments were done on mice expressing the *Drd1-tdTomato* reporter as per our previously published methods.^19^ Briefly, striatal slices from P14-P18 mice were collected and recorded in nominal ACSF. Expression of tdTomato was used to identify the D1-SPNs. Current clamp recordings at incremental current steps (500 ms duration) were applied at resting potential to measure number of action potentials. In the voltage clamp, a –10 mV voltage (500 ms duration) was applied to measure input resistance and normalized cell conductance at –85, -65, and –55 mV holding potentials. IV-plots were generated in voltage clamp conditions using a multi-step protocol (500 ms duration) ranging from –30 to –120 mV voltages at -10mV steps. Current measured was average from a 200 ms window at the end of each step.

### RNA Extraction and quantitative real time PCR (RT-qPCR)

We adapted a previously published protocol to obtain single cell suspensions.^18^ Briefly, mice were anesthetized and sacrificed at P14 and brains were removed and placed in ice-cold ACSF bubbled with 95% O2 and 5% CO2. 500µM coronal sections were made in ACSF and transferred to another chamber filled with ACSF. From 3 striatal sections, 6 punches of striatal tissue were collected (2 per hemisphere). Papain was added and samples were incubated at 37C for 30 minutes. Samples were then triturated using glass pipette. Samples were centrifuged and then resuspended in ACSF. Fluorescence-activated Cell Sorting (FACS) was used to sort D1-SPNs (marked by td-Tomato) from the remaining striatal neurons. RNA from FACS sorted cells was isolated using miRNAeasy kit guidelines. RNA was converted to cDNA using recommended guidelines from SSIII Superscript Kit (Invitrogen) and RT-qPCR was performed using the CFX384 Real-Time System (Bio-Rad). Expression of *Foxp1* and *Kcnk2* was measured from both the td- Tomato positive (D1-SPNs) and td-Tomato negative (non D1-SPN striatal cells) fractions and normalized to expression of 18S.

### AAV-mediated injection of *Foxp1*

A pAAV-hSYN-Foxp1-T2A-eGFP (or control AAV9-hSYN1-eGFP construct expressing only EGFP) was injected into the striatum of mice at postnatal day 1 in litters that were expected to include Foxp1/2^D1^ mice. The titers for each construct were 3.4 X 10^12^ and 2.7 X 10^12^ GC/mL respectively. Pups were anesthetized on ice and then bilateral injections were performed using a beveled glass injection pipette and a Nanoject^TM^ injector (Drummond Scientific). They were then allowed to recover on a heating pad before being returned to their dams. Mice were genotyped before P14 to confirm if they were controls or Foxp1/2^D1^ and whether they carried td-Tomato. Control and rescue virus injected wild type and double knockout mice that were td- Tomato positive were used for electrophysiology experiments. Neurons co-expressing td-Tomato (marking D1-SPNs) and EGFP (marking viral infected neurons) were recorded from as previously described. Any mice not used for electrophysiology were utilized for behavioral experiments (nest building, open field, and rotarod) when they were older than P56.

## Excel File 1

**Supplemental Table 1**: Differentially expressed genes in knockout conditions

Differentially expressed genes in the D1-SPNs for all three knockout conditions. Data provided for juvenile (P9) and adult (P56) as well as cortical neurons from the adult. Striatal DEGs overlapping with the SFARI database of ASD risk genes identified. GO terms associated with each striatal DEG also indicated in separate tabs. Only filtered DEGs (adjusted p-value < 0.05, |logFC| > 0.25) listed.

## Excel File 2

**Supplemental Table 2**: Differentially accessible regions in knockout conditions.

Differentially accessible regions in the D1-SPNs for all three knockout conditions. Genes with DARs enriched for FOX motifs also identified.

